# SpecImmune accurately genotypes diverse immune-related gene families using long-read data

**DOI:** 10.1101/2025.02.04.636381

**Authors:** Shuai Wang, Xuedong Wang, Mengyao Wang, Qian Zhou, Shuai Cheng Li

## Abstract

Polymorphic immune-related genes (HLA, KIR, IG, TCR, and CYP) exhibit significant complexity due to their extensive heterozygosity and inter-loci homology, necessitating specific methods for accurate characterization. We present SpecImmune, the first comprehensive tool leveraging long-read sequencing data to resolve the full spectrum of these immune-related genes. The method adopts an iterative graph-based algorithm for haplotype reconstruction. We validated SpecImmune across 1,019 samples from the 1kGP ONT cohort, 42 PacBio CLR and 9 PacBio HiFi samples from the HGSVC project, and 47 PacBio HiFi and 37 ONT samples from the HPRC project. SpecImmune achieved an accuracy of 98% in HLA typing, which represents a 12% improvement over both SpecHLA and HLA*LA. SpecImmune is the initial method to type multiple CYP loci, as well as the foremost approach to allow precise KIR and germline IG/TCR typing using long reads. Comprehensive genotyping of these loci by SpecImmune unveils a new observation of substantial linkage disequilibrium among HLA, KIR, and CYP loci. The proteins derived from these loci exhibit strong binding affinities, which suggest the origin of the marked linkage disequilibrium. Further, SpecImmune unveils a novel finding of elevated IG/TCR heterozygosity in African populations. Additionally, SpecImmune facilitates the detection of *de novo* mutations and enables allele-specific drug recommendations.

## Introduction

The human immune system encompasses various gene families that are fundamental to protecting against pathogens, including the Human Leukocyte Antigen (HLA), Killer Immunoglobulin-like Receptor (KIR), Immunoglobulin (IG), T cell receptor (TCR), and cytochrome P450 (CYP) families. The HLA locus, located on the short arm of chromosome 6 (6p21.31), is the most gene-dense and polymorphic region in the human genome^1^. Variants frequently occur in the antigen-binding sites of these molecules, which are critical for their interactions with TCRs and KIRs, enabling the immune system to recognize a wide array of antigens. The KIR gene family on chromosome 19q13.4 also exhibits significant allelic polymorphism, consisting of 15 genes and two pseudogenes. Particularly, exon 5 of KIR genes, responsible for encoding the immunoglobulin-like domains crucial for binding to HLA molecules, exhibits elevated levels of nonsynonymous mutations^2^. Inhibitory KIR receptors are essential for maintaining self-tolerance by recognizing self-HLA class I molecules and inhibiting natural killer (NK) cell activity^3^. In contrast, activating KIR receptors detect altered-self or non-self antigens, triggering NK cell-mediated immune responses^4^. The stochastic expression of activating and inhibitory KIR receptors results in a highly diverse NK cell population^4^. The IG genes encode B cell receptors (BCRs), which recognize antigens and initiate antibody production. The TCR genes encode T cell receptors, enabling T cells to recognize antigen-HLA complexes presented by antigen-presenting cells^5^. Both IG and TCR genes undergo V(D)J recombination during lymphocyte development, where variable (V), diversity (D), and joining (J) gene segments are rearranged to generate an extensive repertoire of receptors^5,6^. This diversity is essential for enabling the immune system to recognize and respond to various antigens. The CYP enzyme family, typically associated with drug metabolism, is also essential in immune regulation. CYPs metabolize various compounds that can influence immune responses, including drugs, toxins, and endogenous molecules^7,8^. In cancer patients, CYP activity is often reduced, and the administration of anti-PD-1 antibodies—a form of immune checkpoint inhibitor therapy—can further impair CYP-mediated metabolism^9,10^. This reduction in CYP activity may affect the immune system’s ability to eliminate tumors by altering the pharmacokinetics of immunotherapies and other treatments. Interestingly, studies suggest that enhancing CYP activity could improve the effectiveness of cancer immunotherapies by optimizing drug metabolism and reducing immune suppression^9,10^.

Different immune-related gene families have a complex interplay relationship. The combined diversity of IG, TCR, and HLA molecules forms the foundation of both humoral and cellular immune responses^11^. HLA class I molecules present intracellular antigens to CD8+ T cells, while HLA class II molecules present extracellular antigens to CD4+ T cells^12^. B cells internalize antigen-BCR complexes, process the antigens, and present these as antigen-HLA complexes to T helper cells. The recognition of these complexes by BCRs and TCRs is driven by V(D)J recombination during lymphocyte development^13–15^. Concurrently, NK cells rely on diverse KIRs to interact with HLA class I molecules, which regulate NK cell-mediated cytotoxicity. The specific interactions between KIRs and TCRs with the peptide-HLA class I complex depend highly on the peptide sequence, emphasizing the importance of peptide specificity in immune recognition^16^. The interplay between KIR and HLA gene families has also been shown to influence susceptibility to autoimmune diseases and to impact the success of hematopoietic stem cell transplantation^12^. The impact of CYPs on immune function through drug metabolism and immune modulation underscores the intricate interplay between CYPs and other immune-related gene families in shaping immune responses and treatment outcomes^10,17^. Accurate typing of these gene families would lay a solid foundation for understanding the complex interplay of these genes and comprehending immune function and disease susceptibility.

Many computational methods have been developed for typing these immune-related genes based on short-read sequencing data. HLA typing methods, including Optitype^18^, Polysolver^19^, PHLAT^20^, HLA-VBSeq^21^, HLA-HD^22^, T1K^23^, and others, are designed to ascertain a pair of alleles for each HLA locus. Alternatively, methods like HLA*PRG^24^, HLA*LA^25^, and arcasHLA^26^ utilize reference graphs derived from the HLA database to select suitable HLA types. Additional methods such as Kourami^27^, HISAT-genotype^28^, and SpecHLA^29^ focus on reconstructing HLA sequences through assembly or variant phasing. KIR typing is addressed by methods like KIR*IMP^30^, KPI^31^, PING^32^, and T1K^23^, which analyze KIR genes at the gene-level or identify KIR types from targeted amplified data or whole-genome sequencing (WGS) data. Somatic IG/TCR types identification is pursued by methods like MiXCR^33^, BASIC^34^, and IgCaller^35^. While germline variants are linked to TCR function and TCR repertoire characteristics^36^, the current approach to identifying germline IG/TCR variants faces limitations. ImmunoTyper-SR provides a solution by facilitating germline IGHV genotyping through short-read WGS data^36^. *CYP2D6* allele typing from short-read targeted or WGS data is tackled by methods such as Cypiripi^37^, Astrolabe (previously Constellation)^38^, Aldy^39^, Stargazer^40^, and Cyrius^41^.

Despite advancements in computational algorithms, the short read length of traditional sequencing technologies limits their accuracy and resolution in immune-related gene typing. Genotyping these genes presents several challenges, including high polymorphism, structural complexity, and significant sequence similarity across loci within the same gene family, which hinder accurate read mapping, assembly, and variant phasing^23,29^. For instance, DNA sequence polymorphisms separated by distances longer than the read length lack reliable phasing evidence, leading to ambiguities in typing. Additionally, gene fusions and structural variants (SVs) at immune-related loci further complicate the analysis, as short reads are inadequate for their reliable identification^23,42^. The emergence of long-read sequencing technologies has provided a solution to these challenges, enabling high-resolution characterization of complex immune-related genes^43^. Pacific BioSciences (PacBio) SMRT sequencing generates continuous long reads, including continuous long reads (CLR) and highly accurate circular consensus sequencing (CCS) reads, with the latter achieving lengths of 10–25 Kb and accuracy exceeding 99.5%^44^. Oxford Nanopore Technology (ONT), capable of producing ultra-long reads (>1 Mb)^45^, further enhances the ability to span immune-related loci, providing long-range phase information and enabling rapid typing for clinical applications. These technologies have been successfully applied to elucidate immune-related genes, including HLA^45,46^, KIR^47^, IG and TCR^48,49^, and *CYP2D6*^42^, through strategies such as targeted amplicon sequencing, multiplex PCR, and *de novo* assembly^50^. To address the computational demands of analyzing long-read data, several bioinformatics tools have been developed. Early efforts relied on in-house scripts, which, while innovative, often lacked benchmark validation and robustness^43,46^. More recent software packages such as GenDX NGSengine (https://www.gendx.com/product_line/ngsengine/), HLA*LA^25^, and SpecHLA^29^ have improved HLA typing but face limitations such as high computational costs and reliance on specific sequencing platforms. Tools like pbaa and pangu focus exclusively on PacBio HiFi data, restricting broader applicability^42^. Similarly, current methods for CYP typing primarily target the *CYP2D6* locus and fail to explore interactions across gene families. These challenges underscore the need for versatile and efficient computational methods that can handle diverse immune-related gene families and leverage the full potential of long-read sequencing data.

To address this gap, we have developed an open-source software package called SpecImmune to facilitate rapid and precise typing of diverse immune-related genes from WGS or targeted long-read data. SpecImmune selects a pair of alleles for each locus from the database that optimally corresponds to the sequencing reads, subsequently mapping the reads onto this allele pair to identify variants. Next, SpecImmune applies an iterative haplotype reconstruction algorithm to disentangle the conjugate graph structure, generating linear personalized haplotypes by traversing the graph paths. Allele-specific reads are then realigned to the personalized haplotypes, and the processes of variant calling, segmentation, and maximum matching are iterated until no additional variants are identified. SpecImmune demonstrated superior accuracy in typing HLA, KIR, IG, TCR, and CYP loci in five long-read sequencing datasets: 1,019 ONT sequencing samples from the 1000 Genomes Project (1kGP) cohort^51^, 47 PacBio HiFi samples and 37 ONT samples from the Human Pangenome Reference Consortium (HPRC) project^52^, 42 PacBio CLR samples, and 9 PacBio HiFi samples from the Human Genome Structural Variation Consortium (HGSVC) project^53^. By harnessing SpecImmune’s comprehensive typing within the 1kGP ONT cohort, we discovered a pronounced linkage disequilibrium (LD) among HLA class I and II, KIR, and CYP loci. The proteins of these linked loci demonstrate robust binding affinities, showcasing profound structural compatibility and intermolecular interactions. Moreover, SpecImmune reveals a novel discovery of heightened IG/TCR heterozygosity in African populations while also facilitating the detection of *de novo* mutations (DNMs) and offering allele-based drug recommendations. Our findings underscore the reliability of SpecImmune in typing multiple immune-related gene families, providing valuable insights for immune research.

## Results

### Overview of SpecImmune

Given long-read WGS data, SpecImmune initially extracts reads from specific regions of interest (HLA, KIR, IG, TCR, CYP) aligned to the hg38 reference (Figure 1a-b). This step is skipped for targeted amplicon sequencing data. Subsequently, SpecImmune further bins extracted reads to their respective gene loci by aligning them to the allele database encompassing all alleles within the region of interest (Figure 1c). Next, SpecImmune aligns the binned reads to all alleles at each respective gene locus. These alignments for every read are preserved for subsequent analysis. For each gene locus, SpecImmune records the alignment identity and the number of matching DNA bases for each read aligned to each allele (Figure 1d). It then evaluates all possible allele pairs by computing the total alignment identity and the number of matching bases from all reads for the locus. The allele pair that maximizes alignment identity and matching base count across all binned reads is selected as the best-matched for the locus (Figure 1e).

**Figure 1.**
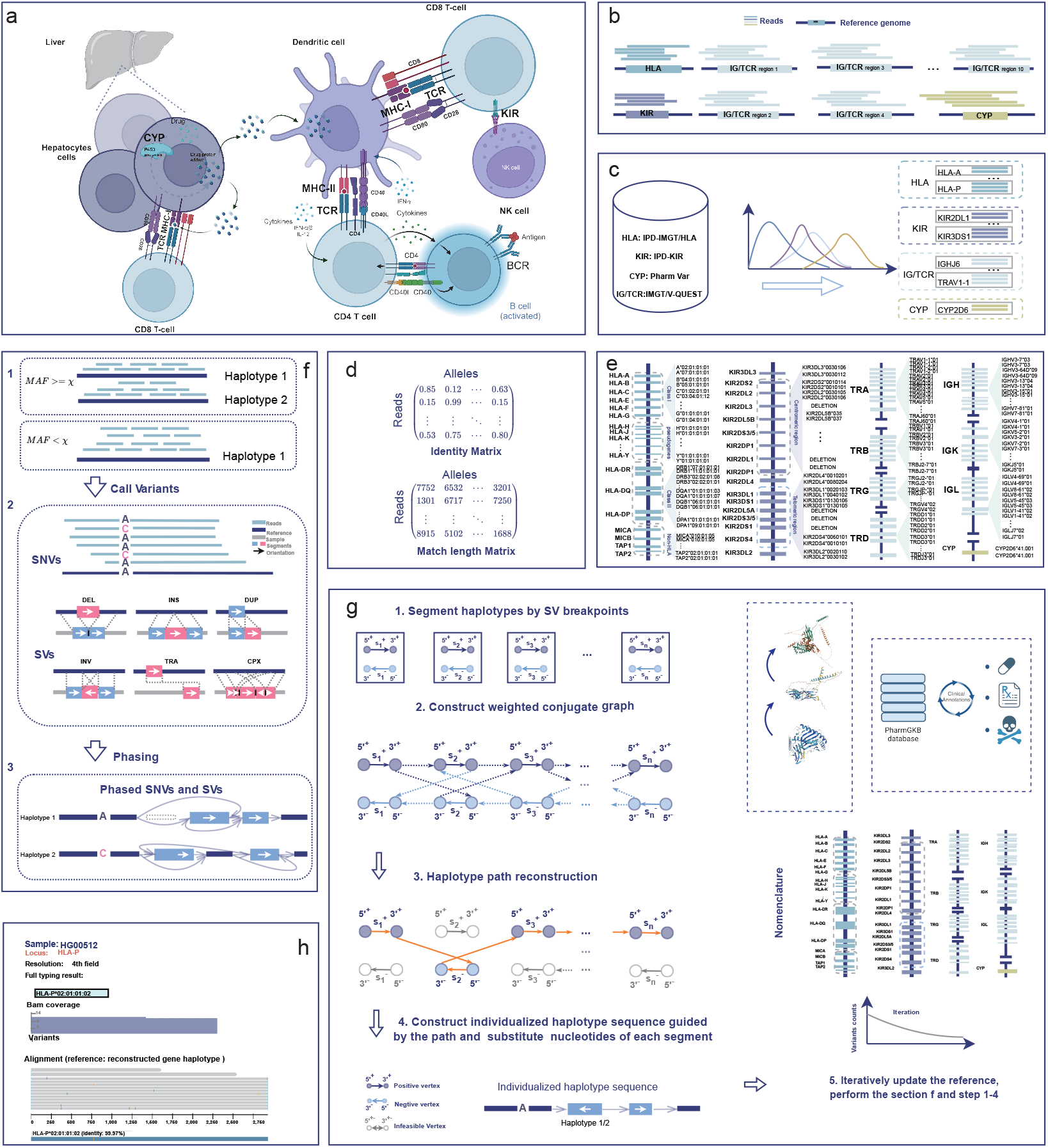
Workflow of SpecImmune. (a) Interactions between HLA, KIR, IG, TCR, and CYP in immunity and metabolism. (b) WGS reads are mapped to the hg38 reference genome, and SpecImmune extracts reads from immune-related gene regions. (c) Reads are binned to gene loci by mapping to the allele database. (d) Total alignment identity and matched base number are estimated for every pair of distinct alleles. (e) Best allele pair is identified, and reads are assigned to alleles. (f) Allele-specific reads are mapped to alleles to construct the haplotype allele, variants calling, and phasing. (g) Iteratively construct personalized haplotype sequence using *haplotype reconstruction* algorithm^58,59^. (h) Visualization of an example results. It shows reads aligned to the reconstructed allele sequence (middle gray bars), alignment coverage (top gray blue distribution), and the typed allele (bottom blue bar) aligned to the reconstructed gene haplotype sequence.

SpecImmune subsequently reconstructs personalized haplotype sequences by leveraging reads aligned to the best-matched allele pair. The process begins with variant detection using Longshot v1.0.0^54^ for SNVs and Sniffles v2.2^55^ for SVs. Joint phasing of SNVs and SVs is then performed using Whatshap v2.3^56^ and Longphase v1.7.3^57^. Based on the identified variants, linear haplotype sequences are reconstructed by applying the haplotype reconstruction algorithm ^58,59^ to resolve non-linear connections. This process is iteratively refined, with haplotypes updated until no new variants are detected. A sliding-window approach is then used to mask low-depth regions, ensuring the generation of high-quality personalized diploid haplotype sequences (Figure 1f-g). Finally, SpecImmune aligns each reconstructed sequence to the allele database and identifies the most matching alleles for the nomenclature of the sequence. Additionally, the allele-specific reads are aligned to the most matched alleles, and the alignments are visualized in an IGV-like figure (Figure 1h). This visualization allows users to easily observe novel variants and the confidence of typing results. Figure 1h illustrates an example of the visualization reports. SpecImmune is the most comprehensive method to type immune-related genes from various sequencing settings (Table 1). The nomenclature schemes for HLA/CYP/KIR alleles are depicted in Supplementary Figure S1. The summary of SpecImmune typing results for the sample HG00377 from the 1kGP is depicted in Supplementary Figure S2.

**Table 1.** Functionalities of state-of-the-art tools in typing immune-related genes from long-read data. √ indicates the software has the feature.

| Methods | HiFi | CLR | ONT | Amplicon | WGS | RNA-Seq | HLA | KIR | IG | TCR | CYP |
| --- | --- | --- | --- | --- | --- | --- | --- | --- | --- | --- | --- |
| SpecImmune | ✓ | ✓ | ✓ | ✓ | ✓ | ✓ | ✓ | ✓ | ✓ | ✓ | ✓ |
| SpecHLA<sup>29</sup> | ✓ | ✓ | ✓ | ✓ | ✓ |  | ✓ |  |  |  |  |
| HLA*LA<sup>25</sup> | ✓ | ✓ | ✓ | ✓ | ✓ |  | ✓ |  |  |  |  |
| NGSEngine | ✓ | ✓ | ✓ | ✓ | ✓ | ✓ | ✓ |  |  |  |  |
| pbaa | ✓ |  |  | ✓ |  |  | ✓ | ✓ |  |  |  |
| PLASTER<sup>42</sup> | ✓ |  |  | ✓ |  |  |  |  |  |  | ✓ |
| pangu | ✓ |  |  | ✓ |  |  |  |  |  |  | ✓ |

Furthermore, SpecImmune is computationally efficient in typing these immune-related gene families (Supplementary Note S1 and Figure S3-4). With its low computational resource requirements, SpecImmune can be run on a personal computer, making it convenient for clinical use. The following sections showcase its typing accuracy and practical application in real-world data analysis.

### SpecImmune is accurate for four-field HLA typing using long-read data

SpecImmune enables accurate four-field HLA typing across various long-read sequencing protocols, supports RNA-seq data, and expands coverage to a larger set of HLA genes compared to existing methods. It identifies 39 MHC-region genes from the IMGT database, including 6 HLA class I genes, 12 HLA class I pseudogenes, 17 HLA class II genes, and 4 non-HLA genes. To validate SpecImmune for HLA typing, we compared SpecImmune with SpecHLA and HLA*LA in five real long-read datasets (Table 2). The GenDX NGSengine software is excluded as it is not freely available. The benchmark datasets include 47 PacBio HiFi samples and 37 ONT samples from the HPRC project, 42 PacBio CLR samples and 9 PacBio HiFi samples from the HGSVC project, and 1,019 ONT sequencing samples from the 1kGP cohort.

**Table 2.** Summary of benchmark datasets in validation of SpecImmune. Due to varying read coverage among gene families in each sample, and the unreliable inference of ground truth for certain gene families in specific samples, the number of samples utilized differs between gene families.

| Dataset | Sequencing protocol | Sequencing strategy | DNA/RNA | Phased assembly available | No. of samples |
| --- | --- | --- | --- | --- | --- |
| 1kGP | ONT | WGS | DNA | No | 1,019 |
| HPRC HiFi | PacBio HiFi | WGS | DNA | Yes | 47 |
| HPRC ONT | ONT | WGS | DNA | Yes | 37 |
| HGSVC HiFi | PacBio HiFi | WGS | DNA | Yes | 9 |
| HGSVC Pair-end | NGS | WGS | DNA | Yes | 32 |
| HGSVC CLR | PacBio CLR | WGS | DNA | Yes | 42 |
| HGSVC Pair-end | NGS | WGS | DNA | Yes | 32 |
| *CYP2D6* Amplicon | PacBio HiFi | Targeted | DNA | No | 10 |
| RNA 1 | Iso-seq | RNA-Seq | RNA | No | 3 |
| RNA 2 | MAS-seq | RNA-Seq | RNA | No | 3 |
| RNA 3 | ONT direct RNA | RNA-Seq | RNA | No | 3 |
| Total | - | - | - | - | 1,173 |

The specific command parameters for SpecHLA, HLA*LA, and SpecImmune are outlined in Supplementary Table S1.

SpecImmune consistently demonstrated the highest accuracy at the four-field HLA typing, outperforming the other tools in every instance in HPRC HiFi, HPRC ONT, HGSVC HiFi, and HGSVC CLR datasets. In the HPRC HiFi dataset, SpecImmune achieved an average typing accuracy of 98% (2,361/2,420), outperforming SpecHLA at 87% (626/720) and HLA*LA at 87% (1,145/1,320). In the HPRC ONT dataset, SpecImmune reached 98% (1,865/1,946) accuracy, while SpecHLA and HLA*LA achieved 69% (386/592) and 86% (910/1,058), respectively. For the HGSVC CLR dataset, SpecImmune recorded an accuracy of 95% (1,101/1,154), with SpecHLA and HLA*LA reaching 86% (257/300) and 16% (101/570), respectively. In the HGSVC HiFi dataset, SpecImmune attained 98% (524/536) accuracy, compared to 87% (123/142) for SpecHLA and 88% (231/262) for HLA*LA (Figure 2a-d). The number of processed gene loci varied among the tools, and we calculated the accuracy based on all processed gene loci for each tool. Moreover, we categorized the HLA genes into four classes: Class I, Class I pseudogenes, Class II, and Non-HLA. Subsequently, we computed the accuracy for each class individually. In these four high-quality datasets, SpecImmune achieved typing accuracies of 99% (1,007/1,010), 96% (1,088/1,132), 93% (2,034/2,182), and 82% (551/674) across the four gene classes on average. In comparison, SpecHLA reached 88% (442/504) accuracy for class I genes and 76% (640/838) for class II genes. HLA*LA attained accuracies of 76% (763/1,010), 65% (176/272), and 67% (733/1086) for class I, class I pseudogenes, and class II genes, respectively (Figure 2a-d). For each HLA gene class, SpecImmune demonstrated superior accuracy compared to the other two software tools. Additionally, SpecImmune consistently achieved higher accuracy for each HLA gene than the other two methods (supplementary Figure S5-8).

**Figure 2.**
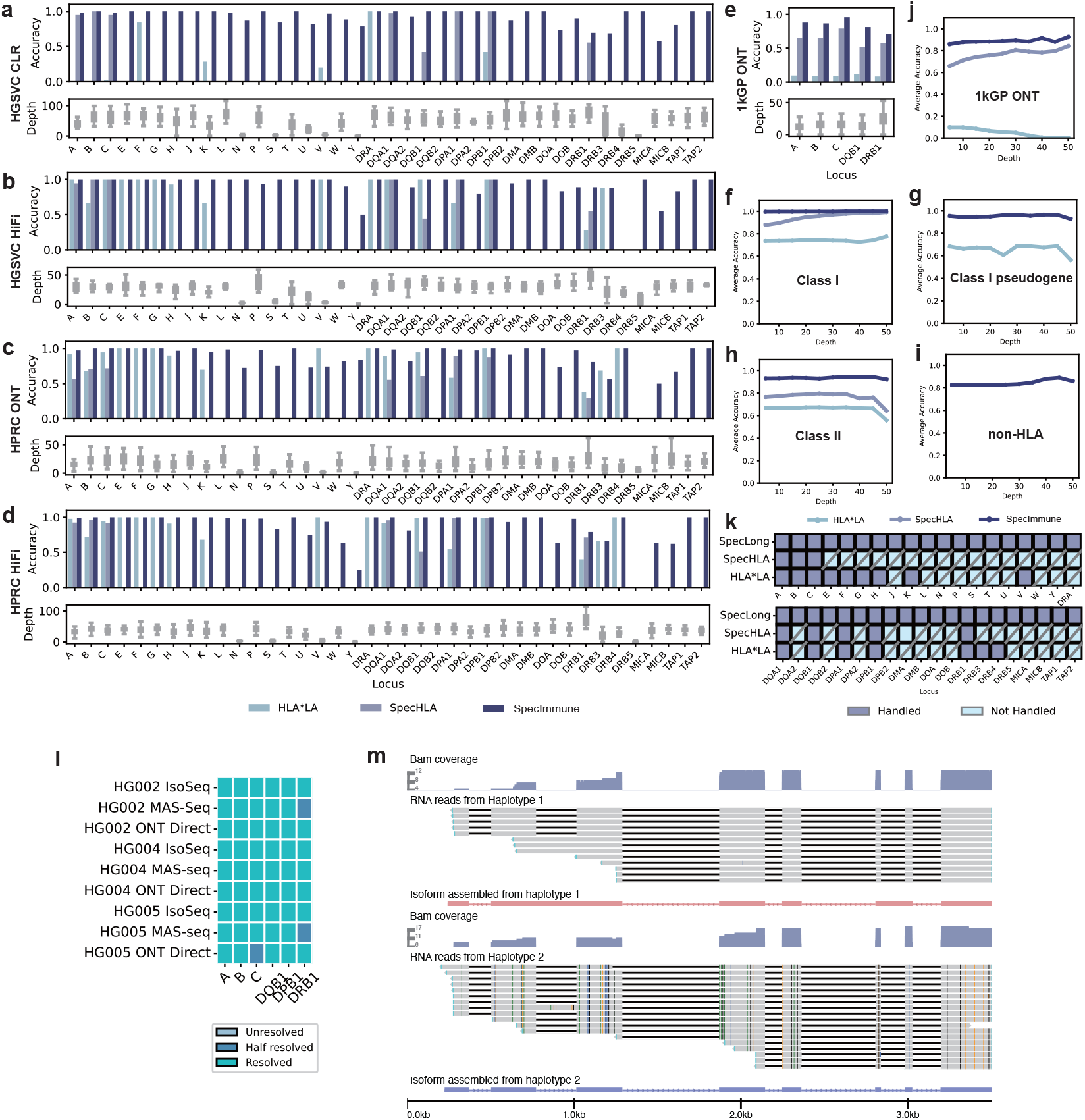
Evaluation of SpecImmune for typing HLA genes. (a–d) Accuracy comparisons among SpecHLA, HLA*LA, and SpecImmune, along with depth distributions across HLA loci for the HGSVC CLR (a), HGSVC HiFi (b), HPRC ONT (c), and HPRC HiFi (d) datasets, respectively. (e) Accuracy comparisons and depth distributions across the *HLA-A, HLA-B, HLA-C, HLA-DQB1*, and *HLA-DRB1* loci within the 1kGP ONT dataset. (f–i) Average performance of SpecHLA, HLA*LA, and SpecImmune across class I (f), class I pseudogene (g), class II (h), and non-HLA (i) classes as sequencing depth increases, aggregated over four different datasets. (j) Average performance of the three methods in the 1kGP ONT dataset with increasing sequencing depth. (k) Ability to handle HLA genes among the three methods.(l) Performance of SpecImmune across various long-read RNA-Seq technologies. “Resolved” indicates that both haplotypes of the locus were successfully typed, “Half resolved” indicates that only one of the two haplotypes was successfully typed, and “Unsolved” indicates that neither haplotype was successfully typed. (m) Phased long RNA-Seq reads alongside allele-specific isoform assemblies for *HLA-A* in the HG002 sample. The first track shows the reads coverage of haplotype 1, the second track shows the alignment of haplotype 1 reads, and the third track shows the assembled RNA isoform expressed from haplotype 1. The fourth track shows the reads coverage of haplotype 2, the fifth track shows the alignment of haplotype 2 reads, and the sixth track shows the assembled RNA isoform expressed from haplotype 2.

Further, among the three tools, SpecImmune again demonstrated the highest accuracy in the intermediate-depth 1kGP ONT dataset. In this dataset, our evaluation focused on two-field typing for *HLA-A*, -B, -C, -DQB1, and -DRB1 genes, as these five genes were the only ones with available reference typing results (Methods). We filtered out low-quality alignments and genes with an average depth below 10, given the generally low sequencing depth in the MHC region for the 1kGP ONT data. SpecImmune achieved accuracies of 94% (1,134/1,208), 89% (1,402/1,568), 98% (1,561/1,590), 85% (1,209/1,422), and 73% (1,348/1,858) across these five genes, whereas SpecHLA reached 82% (996/1,208), 72% (1,133/1,568), 85% (1,357/1,590), 57% (812/1,422), and 59% (1,100/1,858), respectively. HLA*LA showed accuracies of 10% (124/1,208), 9% (137/1,568), 10% (152/1,590), 11% (160/1,422), and 9% (158/1,858) (Figure 2e). SpecImmune’s accuracy was the highest for each gene. HLA*LA’s performance is not higher than 11% for each gene, indicating its poor performance in this ONT dataset.

SpecImmune demonstrated robust typing accuracy across a range of sequencing depths. We evaluated the performance of three software packages at various sequencing depth cutoffs (5x, 10x, 15x, 20x, 25x, 30x, 35x, 40x, 45x, 50x), excluding loci with depths below the specified cutoff at each level. Accuracy was calculated across all samples from four datasets. In the HPRC HiFi, HPRC ONT, HGSVC HiFi, and HGSVC ONT datasets, SpecImmune consistently achieved the highest accuracy across all depth cutoffs (Figure 2f-i). Similarly, in the 1kGP ONT dataset, SpecImmune maintained the highest accuracy at every depth cutoff (Figure 2j). Furthermore, SpecImmune’s accuracy steadily increased as the sequencing depth cutoff rose.

Moreover, SpecImmune is the most comprehensive long-read-based tool capable of handling the majority of HLA genes and can support RNA-seq data. SpecImmune supports HLA typing from a variety of long-read RNA-seq technologies, including PacBio Iso-Seq, ONT 2D cDNA-seq, and Direct RNA-seq, enabling allele-specific isoform assembly and quantification. We validated its performance using nine datasets from the GIAB project^60^, achieving an average accuracy of 97% (105/108) at the G group resolution (Figure 2l). Figure 2m illustrates that long-read RNA-seq reads can span all the exons of *HLA-A* and SpecImmune accurately assembles the isoform. SpecImmune can type much more HLA genes compared to the other tools. As shown in (Figure 2k), SpecImmune can process 39 genes across four gene classes within the MHC region. In contrast, SpecHLA is limited to 8 genes from HLA class I and HLA class II, while HLA*LA is confined to 17 genes across HLA class I, HLA class I pseudogenes, and HLA class II. In all of the HPRC HiFi, HPRC ONT, HGSVC HiFi, and HGSVC CLR datasets, SpecImmune demonstrated the highest average accuracy for the eight genes that all three tools could process and achieved 94% (2,395/2,542)accuracy for the 22 genes that only SpecImmune could handle. To date, SpecImmune is the tool that can handle most of the HLA genes from long-read data.

### SpecImmune effectively determines KIR typing using long-read data

SpecImmune demonstrated accurate and robust typing of KIR genes in four real datasets and one simulated dataset. SpecImmune’s KIR typing performance was analyzed in 9 HGSVC HiFi, 19 HGSVC CLR, 45 HPRC HiFi, and 26 HPRC ONT WGS samples (Table 2). Sequencing reads from these WGS samples were extracted to focus on the KIR region. The published phased assemblies of these samples aided in inferring the reference allele types (Methods). SpecImmune achieved impressive 1-field KIR typing accuracies: 92.9% (589/634) in HPRC HiFi, 94.9% (74/78) in HGSVC HiFi, 93.8% (407/434) in HPRC ONT, and 90.8% (138/152) in HGSVC CLR datasets (Figure 3a-c). The highest accuracy was attained in the HGSVC HiFi dataset. While accuracy slightly declines with higher resolution, SpecImmune maintains high accuracy even at the 3-field level, achieving 90.0% (570/634), 92.3% (72/78), 91.0% (395/434), and 89.5% (136/152) in the respective datasets. SpecImmune exhibited higher accuracy in the HPRC ONT dataset than in the HPRC HiFi dataset; however, the former has limited sequencing reads for the framework gene *KIR3DP1*. Figure 3d-e showcases the allele counts inferred by SpecImmune in each dataset, revealing that the framework genes *KIR3DL3, KIR3DP1, KIR2DL4*, and *KIR3DL2* had more alleles detected compared to other KIR genes, with *KIR3DP1* showing particularly low counts in the HPRC ONT dataset. Additionally, we assessed the proportion of samples with at least five reads in the *KIR3DP1* locus in each dataset, yielding proportions of 88.9%, 100%, 32.4%, and 84.2% in HPRC HiFi, HGSVC HiFi, HPRC ONT, and HGSVC CLR datasets, respectively. This suggests potential lower sequencing sensitivity for *KIR3DP1* with ONT technology. SpecImmune demonstrated the lowest accuracy in the HGSVC CLR dataset. However, increasing the read cutoff significantly enhanced the accuracy of this dataset. At the 3-field level, accuracy in the HGSVC CLR dataset surged to 92.4% with a minimum of 20 reads.

**Figure 3.**
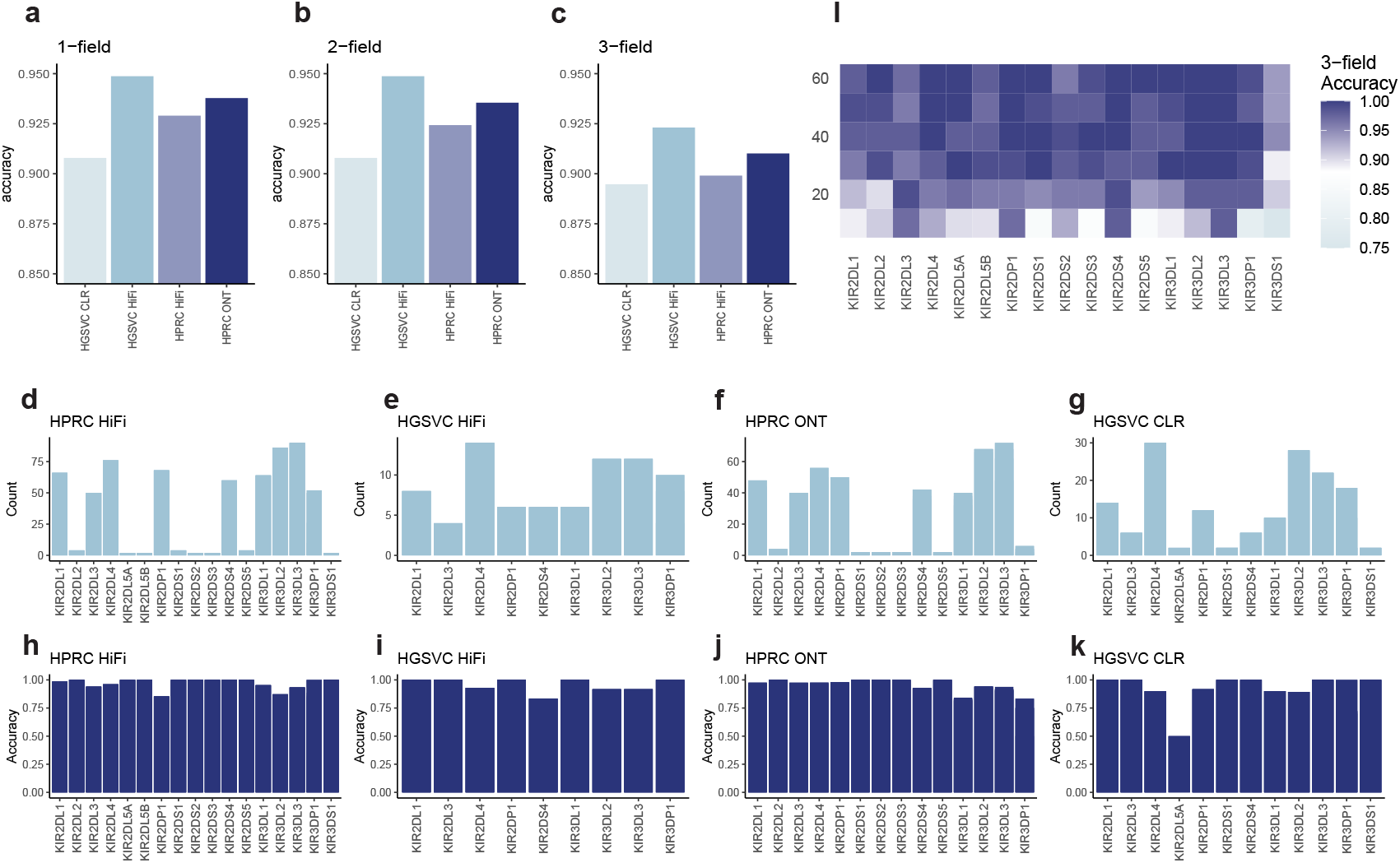
Evaluation of SpecImmune for typing KIR genes. (a-c) SpecImmune’s KIR typing accuracy in four real datasets at 1-field (a), 2-field (b), and 3-field (c) resolutions. (d-g) The count of correctly detected alleles and total detected alleles at each KIR locus in the HPRC HiFi (d), HGSVC HiFi (e), HPRC ONT (f), and HGSVC CLR (g) datasets. (h) Accuracy of 3-field KIR typing at various depth levels in simulated data.

Moreover, simulated data has demonstrated that SpecImmune excels in accurately typing each KIR gene. To carry out this evaluation, we simulated a dataset with sequencing depths ranging from 10x to 60x, with increments of 10x. Each depth setting was replicated 50 times, resulting in 3,000 samples. The data were simulated using PBSIM3^61^, and the sequencing accuracy was set at 90%. At a sequencing depth of 10x, the accuracy for each gene was relatively lower compared to higher depths (Figure 3h). The accuracies among the genes ranged from 75% to 98%, with an average accuracy of 90.0% across all genes at this depth. As the sequencing depth increased, so did the accuracy. For depths exceeding 10x, the accuracy for all genes consistently remained at or above 90%. By the time the sequencing depth reached 60x, the average accuracy for all genes peaked at an impressive 98.9%. The evaluation in simulated data underscores the robustness and reliability of SpecImmune in KIR gene typing tasks.

### SpecImmune accurately performs germline IG/TCR typing using long-read data

Benchmark results demonstrated that SpecImmune accurately infers germline IG and TCR types, and can infer haplotypes across different gene loci. SpecImmune was validated in 9 HGSVC HiFi, 19 HGSVC CLR, 45 HPRC HiFi, and 26 HPRC ONT samples (Table 2). The ground truth references were inferred from phased assemblies of the HGSVC and HPRC individuals (Methods). SpecImmune achieved high IG and TCR typing accuracies of 97.8% (5,440/5,562), 98.5% (11,452/11,622), 98.1% (28,933/29,504), and 95.8% (17,901/18,694) in the HGSVC HiFi, HGSVC CLR, HPRC HiFi, and HPRC ONT datasets, respectively (Figure 4a). SpecImmune exhibited lower accuracy with ONT data than PacBio HiFi and CLR data. As sequencing depth increases, the accuracy in these datasets also improves. With a minimum depth of 20x, in the HGSVC HiFi, HGSVC CLR, HPRC HiFi, and HPRC ONT datasets, SpecImmune achieved IG and TCR typing accuracies of 99.4% (4,703/4,732), 99.1% (11,045/11,144), 99.6% (24,456/24,552), and 99.3% (10,914/10,986), respectively. The accuracy in all four datasets exceeds 99% with a minimum depth of 20x. Furthermore, we classified IG and TCR loci into IGH, IGK, IGL, TRA, TRB, TRD, and TRG categories and counted the average accuracy of each category (Figure 4b-e). Notably, SpecImmune demonstrated relatively lower accuracy for IGH and IGK loci compared to other loci consistently across the four datasets, possibly due to the higher diversity of IGH and IGK loci. With an increase in the sequencing depth, the typing accuracy overall increases, with a more pronounced improvement observed for IGH and IGK loci.

**Figure 4.**
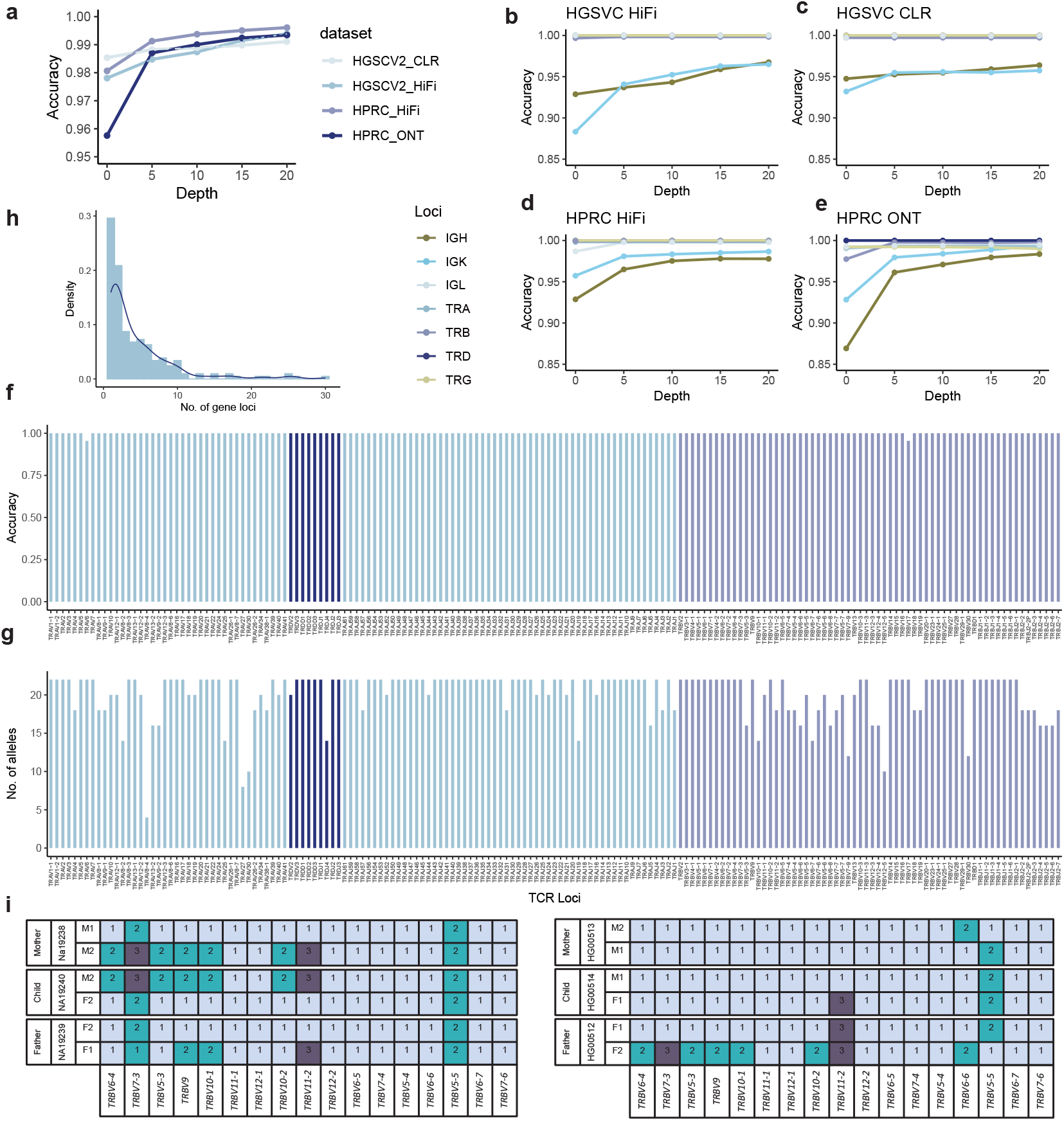
Evaluation of SpecImmune for typing IG/TCR genes. (a) Average typing accuracy of all IG/TCR genes with varying sequencing depth cutoffs. (b-e) Typing accuracy of different IG/TCR gene groups with varied sequencing depth cutoffs in the HGSVC HiFi (b), HGSVC CLR (c), HPRC HiFi (d), and HPRC ONT (e) datasets. (f-g) Accuracy (f) and number (g) of detected alleles at each TCR locus in the targeted TCR HiFi sequencing data. (h) Distribution of heterozygous gene loci count in each phase block. (i) Illustration of the phase block in two family trios. Numbers represent different alleles at each locus.

Furthermore, 11 targeted TCR amplicon HiFi sequencing samples validated SpecImmune’s reliability for germline TCR typing. The reference allele types of the TRA/D and TRB loci were reported by the previous research^49^. The types reported as novel were discarded to ensure a fair evaluation. Also, the loci with sequencing depth lower than two were discarded. There were 3,410 alleles of these TRA/D and TRB loci remaining after the discarding. In these alleles, SpecImmune successfully typed 3,408 alleles, with an accuracy of 99.9% (Figure 4f). Notably, SpecImmune achieved a 100% accuracy for all the loci except for *TRAV5* and *TRBV17*. Figure 4g shows the number of alleles used to evaluate each locus.

Additionally, SpecImmune infers the haplotype across different gene loci accurately. Leveraging the long reads, SpecImmune can group multiple gene loci into the same phase block. In the 9 HGSVC HiFi samples, the number of heterozygous gene loci of each phase block was counted. A phase block constructed by SpecImmune covers 4.5 heterozygous gene loci on average (Figure 4h). The largest phase block covers 30 heterozygous gene loci. Further, the accuracy of the phase blocks was assessed by trio consistency. The 9 HGSVC HiFi samples consist of three family trios. For every pair of adjacent heterozygous gene loci within the same phase block in the child, we verify their trio consistency by confirming the presence of both haplotypes in the parents. In these three trios, 84.6% (115/136) of the adjacent heterozygous gene loci pair are phased in a trio-consistency manner. We illustrate the haplotypes across 17 gene loci inferred by SpecImmune fit trio consistency in two family trios (Figure 4i).

### SpecImmune enables precise CYP typing using long-read data

SpecImmune excels in accurately typing *CYP2D6* over current methods, especially in error-prone CLR and ONT data. Additionally, SpecImmune effectively types other CYP loci where other methods fall short. SpecImmune was compared with pangu for *CYP2D6* typing in 56 1kGP ONT, 32 HPRC HiFi, 25 HPRC ONT, and 10 HiFi amplicon samples (Table 2). The *CYP2D6* truth derives from the GeT-RM project^62^ and previous studies^42^. The method pangu is designed for HiFi data; as there is a lack of computational tools to type *CYP2D6* based on other long reads sequencing protocols, pangu is also utilized as the baseline method in ONT data besides HiFi data. PLASTER is discarded in comparison as it is only designed for the amplicon data^42^.

SpecImmune performs slightly better than pangu for *CYP2D6* typing in HiFi data (Figure 5a). In 32 HPRC HiFi samples, the *CYP2D6* typing accuracy of SpecImmune and pangu were 100% (64/64) and 95.3% (61/64), respectively. As the number of reads varies between these samples, we set a read count cutoff to evaluate the impact of depth on SpecImmune and pangu. The accuracy of SpecImmune is 100% with different read numbers. While increasing the number of reads, the accuracy of pangu increases. With no less than 50 reads, the accuracy of pangu also achieves 100% (20/20). The result shows that both SpecImmune and Pangu are accurate with HiFi data.

**Figure 5.**
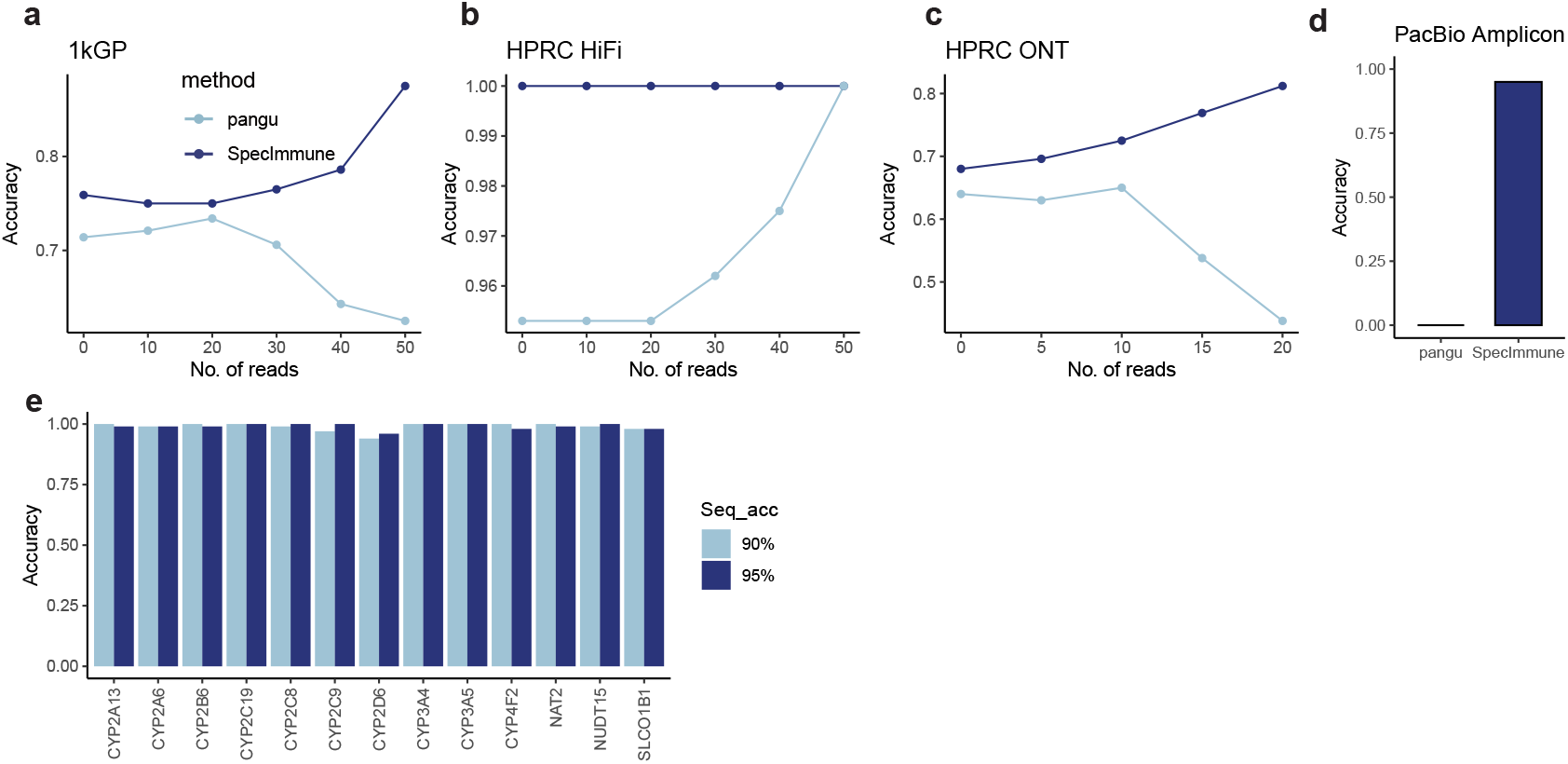
Evaluation of SpecImmune for typing *CYP2D6*. (a-c) Typing accuracy of *CYP2D6* by SpecImmune and pangu in the 1kGP (a), HPRC HiFi (b), and HPRC ONT (c) datasets. (d) Typing accuracy of *CYP2D6* by SpecImmune and pangu in HiFi amplicon data. (e) Accuracy of SpecImmune in typing different CYP loci using simulated data. Bar color indicates sequencing read accuracy at either 90% or 95%.

Notably, in ONT data, SpecImmune consistently outperformed pangu in the inference of *CYP2D6* star alleles, with SpecImmune’s advantage becoming more significant as the number of reads increased. Among the 32 HPRC ONT samples, the accuracies of SpecImmune and pangu were 68% (34/50) and 64% (32/50) respectively. As the read count increased, SpecImmune demonstrated a continuous improvement in accuracy (as shown in Figure 5b). Specifically, with a minimum of 20 reads, SpecImmune achieved an accuracy of 81.2% (13/16). Conversely, the performance of pangu deteriorated with the increasing number of reads, as its accuracy dropped to 43.8% (7/16) with at least 20 reads. In the 56 ONT samples from the 1kGP, the accuracies of SpecImmune and pangu in inferring *CYP2D6* star alleles were 75.9% (85/112) and 71.4% (80/112) respectively. With the increased number of reads, SpecImmune’s accuracy continued to rise (refer to Figure 5c). Notably, with a minimum of 50 reads, SpecImmune achieved an accuracy of 87.5% (7/8). Conversely, the performance of pangu seemed to suffer from an increase in reads, dropping to 64.3% (18/28) with at least 40 reads, and further decreasing to 62.5% (5/8) with at least 50 reads. Both ONT datasets consistently illustrate that SpecImmune is a more reliable choice compared to pangu for analyzing ONT data. The increase in the number of reads consistently benefits SpecImmune while negatively impacting pangu’s performance.

Additionally, SpecImmune is reliable for *CYP2D6* typing in HiFi amplicon data. We run SpecImmune and pangu on 10 HiFi amplicon data. The data only include the reads from the *CYP2D6* region, and it is impossible to infer CNV. Therefore, we ignore the alleles with CNVs when calculating the accuracy in this dataset. The pangu’s results cannot be used in this dataset; it generates over ten haplotypes for each sample. In contrast, SpecImmune achieves an accuracy of 98% (19/20) (Figure 5d). These results showed that SpecImmune is accurate for *CYP2D6* typing in both HiFi and ONT WGS data; it can also handle the amplicon sequencing data.

Furthermore, SpecImmune demonstrates accurate typing capabilities across multiple CYP loci beyond *CYP2D6*. We conducted simulations involving 50 samples with 95% sequencing accuracy and a depth of 100x for all 13 CYP loci. SpecImmune exhibited an overall accuracy of 99.1% (1,288 out of 1,300) in identifying star alleles across these loci (Figure 5e). To gauge SpecImmune’s resilience against sequencing errors, we simulated 50 samples with 90% sequencing accuracy and maintained a depth of 100x for all 13 CYP loci. In this scenario, SpecImmune achieved an accuracy of 98.9% (1,286 out of 1,300), slightly below the accuracy observed with 95% sequencing accuracy. These findings underscore SpecImmune’s reliability in typing various CYP genes from long reads and its robustness in handling sequencing errors.

### SpecImmune analysis uncovers heightened IG/TCR heterozygosity in African populations

Using SpecImmune, we confirmed that African populations exhibit higher IG/TCR heterozygosity and observed strong geographic correlations in HLA, KIR, and CYP allele distributions, consistent with prior studies^41,63,64^. To systematically assess immune-related gene diversity, we analyzed 1,019 1kGP ONT WGS samples from 26 populations, categorized into five superpopulations: 189 from Europe (EUR), 192 from East Asia (EAS), 199 from South Asia (SAS), 275 from Africa (AFR), and 164 from the Americas (AMR) (Figure 6a).

**Figure 6.**
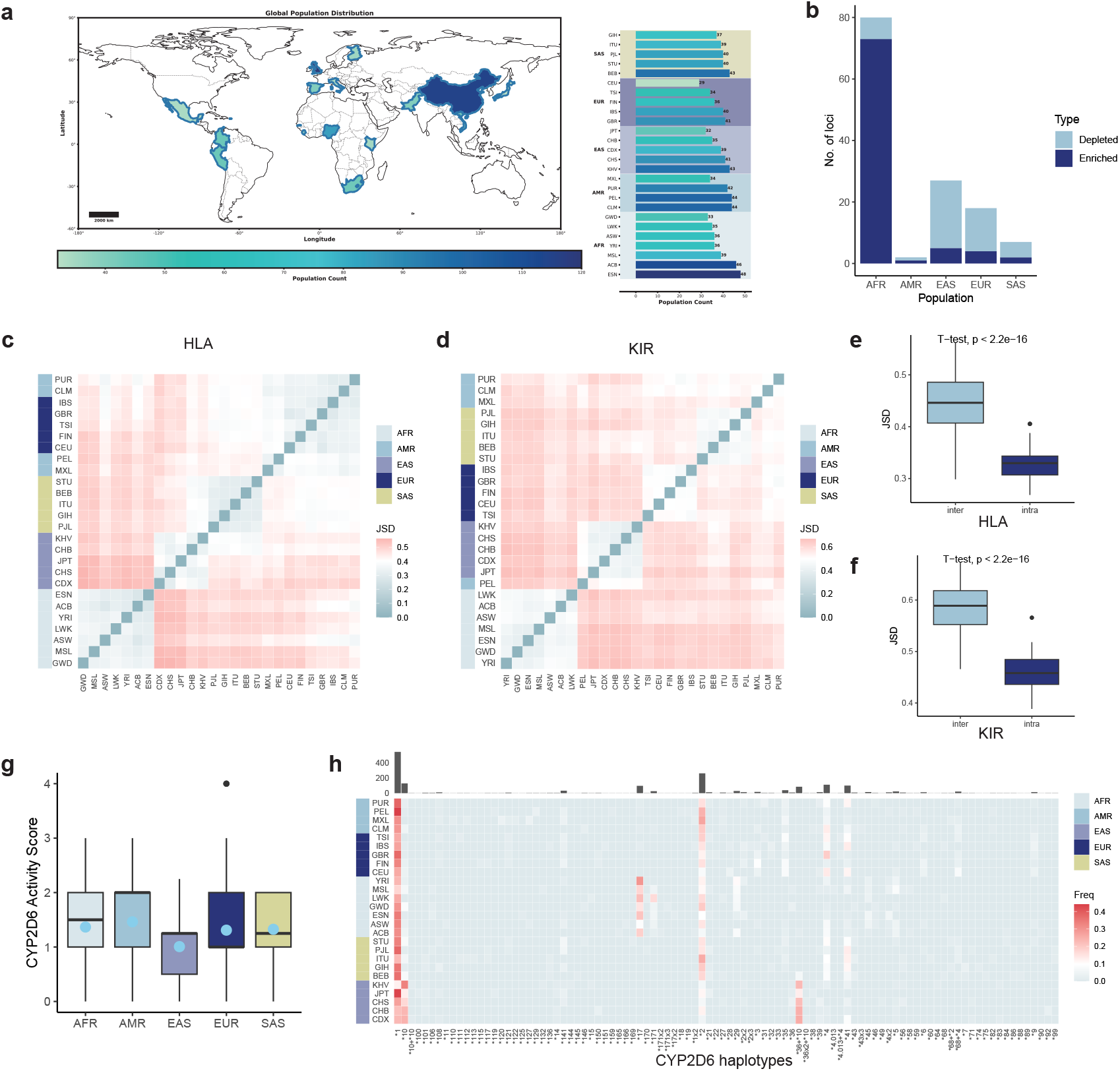
Association between population and immune gene allele distribution in 1kGP. (a) Sample sizes of different populations in the 1kGP dataset. (b) Counts of IG/TCR loci enriched and depleted heterozygous variants across all super populations. (c) Quantification of HLA allele frequency differences using JSD among different populations. (d) Quantification of KIR allele frequency differences using JSD among different populations. (e) Comparison of JSD values within (intra) and between (inter) super populations for HLA alleles. (f) Comparison of JSD values within and between super populations for KIR alleles. (g) Boxplot of *CYP2D6* activity scores in each super population. (h) Heatmap displaying the frequency of each *CYP2D6* allele in each population. The top bar plot illustrates the count of each *CYP2D6* allele detected across all populations.

Our analysis revealed that the African population harbors significantly higher IG/TCR gene heterozygosity compared to other populations. Specifically, among the 80 loci with significant differences in heterozygous variant counts (p.adj < 0.01), 91.2% were enriched in the African population. Notably, loci *IGHV7-81* and *TRGV10* exhibited the most pronounced enrichment. In contrast, other populations showed considerably fewer loci with heterozygous variant enrichment, including EAS (18.5%), EUR (22.2%), SAS (28.6%), and AMR (50%) (Figure 6b). These results underscore the exceptional genetic diversity of African populations, particularly in IG/TCR genes, consistent with their previously established higher overall genetic diversity^51,53^.

As anticipated, HLA, KIR, and CYP allele frequencies demonstrated significant population-specific patterns. Populations within the same superpopulation displayed similar HLA allele frequencies (p < 2.2 × 10^−16^, Figure 6c,e), with KIR alleles showing similar geographic specificity (Figure 6d,f). East Asian populations exhibited significantly lower *CYP2D6* activity scores (p = 3.45 × 10^−12^), with haplotypes \**10* and \**36*+\**10* being highly prevalent in this group, while haplotype \**17* was enriched in African populations (Figure 6g,h). These findings align with prior research linking HLA, KIR, and CYP allele distributions to geographic origins^41,63,64^.

Additionally, we observed distinct variations in HLA and KIR allele frequency and diversity across loci, while significant heterozygosity level disparities were detected among IG/TCR gene loci (Supplementary Note S2-4 and Figure S10-13).

### HLA, KIR, and CYP loci exhibited strong linkage disequilibrium

Strong linkage disequilibrium (LD) was observed among HLA, KIR, and CYP loci. We collected the HLA, KIR, CYP, and IG/TCR alleles for each sample from SpecImmune’s results to estimate the level of LD between every pair of loci within these regions (Methods). Due to the varying numbers of alleles at loci *a* and *b*, in addition to the standard overall LD measure *W*_*n*_, the conditional asymmetric LD (ALD) measures *W*_*a/b*_ and *minALD* were employed^65,66^. The *Wn* measure assumes and enforces symmetry. However, in scenarios involving more than two alleles per locus, unequal allele frequencies at each locus, and differing levels of LD between loci, this assumption of symmetry may not hold^65,66^. The ALD measures were utilized to address this (Methods). To ensure the reliability of the results, a minimum of 500 samples with both loci successfully typed by SpecImmune was required for a gene pair. Following filtering, the LD results were obtained for 35 HLA, 8 KIR, 67 IG, 61 TCR, and 13 CYP loci.

Remarkably, both HLA class I and II loci, along with KIR loci, exhibited strong LD. Loci originating from the same genomic region demonstrated elevated *Wn* values and high *W*_*a/b*_ values with each other, corroborating the established patterns of LD within these regions (Figure 7a-b). As anticipated, the strong LD observed between KIR and HLA class I loci aligns with their documented dynamic co-evolutionary adaptations, driven by environmental pressures and intricate interactions^67–69^. Notably, our findings extend these observations by providing evidence for the co-evolutionary dynamics involving HLA class II loci and KIR. Specifically, we highlight the potential interactions of *HLA-DQB1, HLA-DRB1*, and *HLA-DPB1* with KIR alleles, which may play a critical role in immune response regulation. While earlier studies hinted at the indirect influence of HLA class II molecules on NK cell activity through interactions with *NKp44*^70^, our results substantiate and expand upon these mechanisms, underscoring a broader interplay between HLA class II loci and KIR in immune function.

**Figure 7.**
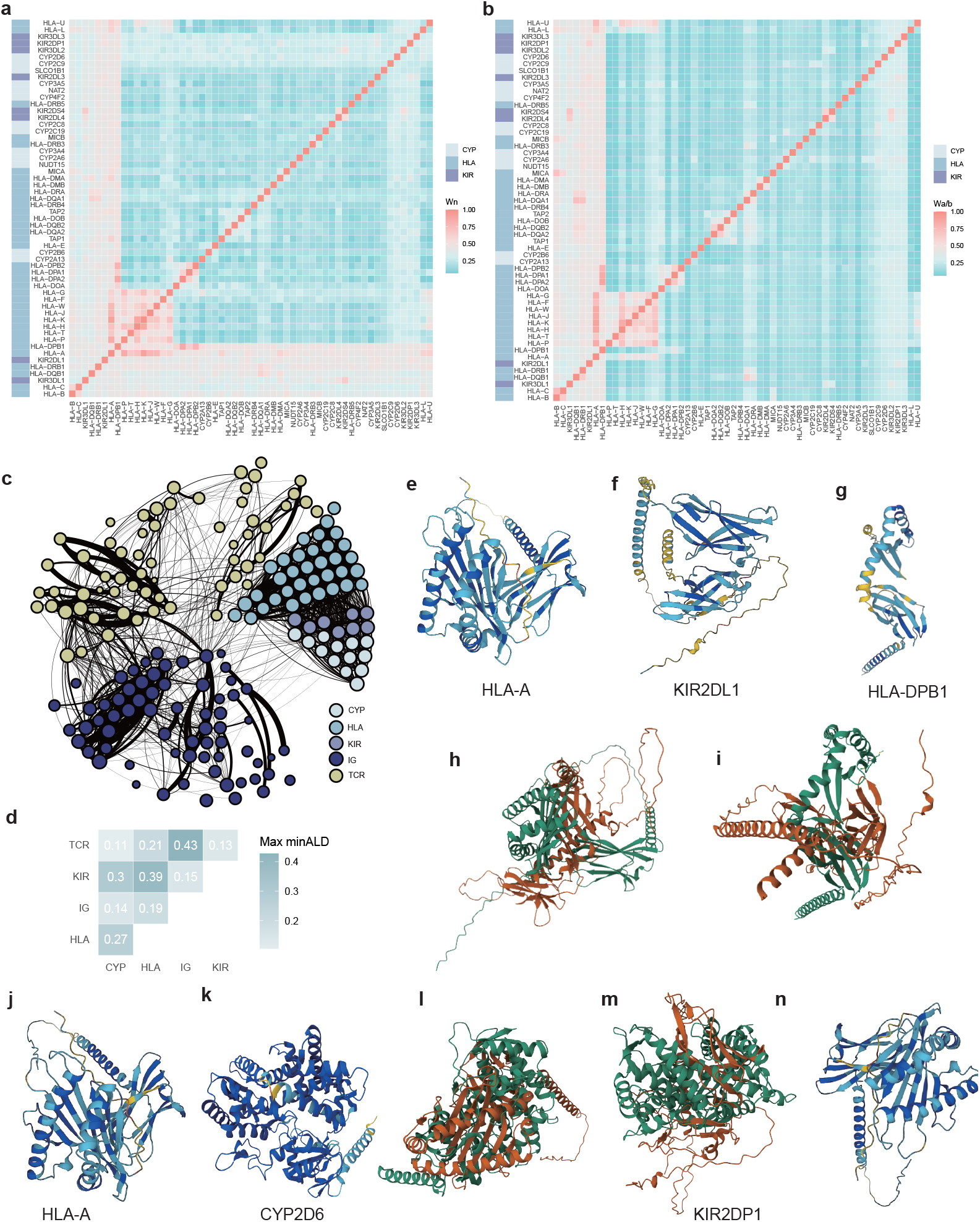
LD between immune-related genes in 1kGP population. (a-b) Heatmap of LD *Wn* (a) and *W*_*a/b*_ (b) values between KIR, HLA, and *CYP2D6* loci. (c) Graph overview of the LD among immune-related genes. Nodes represent specific gene loci, with edge weights reflecting the *minALD* values between them. Node colors indicate gene families, while node sizes correspond to their degrees. Edges with weights below 0.2 are omitted from the visualization. (d) Maximum *minALD* value across immune-related gene regions. (e-n) Predicted 3D protein structure of *HLA-A*\**11:01:01:01* (e), *KIR2DL1*\**0030204* (f), binding affinity of *HLA-A*\**11:01:01:01* and *KIR2DL1*\**0030204* (h), *HLA-DPB1*\**05:01:01:01* (g), binding affinity of *HLA-DPB1*\**05:01:01:01* and *KIR2DL1*\**0030204* (i), *HLA-A*\**01:01:01:01* (j), *CYP2D6*\**1* (k), binding affinity of *HLA-A*\**01:01:01:01* and *CYP2D6*\**1* (l), *KIR2DP1*\**0020119* (m), and binding affinity of *CYP2D6*\**1* and *KIR2DP1*\**0020119* (n).

Furthermore, *CYP2D6* exhibits LD with both KIR and HLA loci (Figure 7c-d). *CYP2D6* is the CYP locus exhibiting the strongest LD with HLA and KIR. Among the KIR and HLA loci, *KIR2DP1* (0.30) demonstrates the highest *minALD* value with *CYP2D6*, followed by *KIR3DL3* (0.274) and *HLA-A* (0.273). Additionally, *CYP2C9* has a high LD value with *KIR2DP1* (0.265). It is known that CYP genes can impact immune function via drug metabolism. This might result in the co-evolution between *CYP2D6*, KIR, and HLA loci^9,10^. Previous studies have shown that drug metabolites, often produced by CYP enzymes, bind to proteins to form complexes. These complexes are processed by antigen-presenting cells (APCs) and presented via MHC II molecules, leading to CD4+ T cell activation. Drugs or metabolites can bind directly to HLA molecules or alter the HLA antigen-binding cleft, activating specific T cells. CD8+ T cells recognize drug-protein complexes presented by MHC I on hepatocytes, leading to their immune destruction. These might lead to the LD between CYP and HLA loci. On the other hand, the LD between CYP and KIR genes may result from their roles in drug metabolism and immune regulation. CYP enzymes generate metabolites that modulate immune responses, while KIR genes regulate NK cell activity in immune surveillance. This genetic linkage may reflect evolutionary pressures favoring specific CYP-KIR combinations to balance drug metabolism and immune function, particularly in drug-induced immune reactions^71^. While specific IG and TCR gene pairs show LD associations; however, IG/TCR regions do not display significant LD with other genetic regions (Figure 7c-d).

Additionally, graph analysis highlighted the significance of *HLA-G, TRBV16* in the immune regulation system.. We constructed an LD graph encompassing all immune-related genes, with genes represented as nodes and edge weights denoting the *minALD* values between respective loci (Figure 7c). The node with the highest degree is *HLA-G*, followed by *CYP2C9, HLA-F, HLA-A*, and *HLA-DOA*. Regarding betweenness centrality, the top-ranking node is *TRBV16*, succeeded by *HLA-DPB2, HLA-DOA, TRBV10-3*, and *HLA-DPA1*. In terms of PageRank score, *TRBV16* holds the highest position, trailed by *HLA-G, HLA-H, HLA-A*, and *HLA-F*.

### Strong binding affinities of HLA, KIR, and CYP proteins underpin marked linkage disequilibrium

HLA, KIR, and CYP proteins exhibit high binding affinities and structural compatibility, suggesting their LD might derive from molecular interactions. To investigate this, we selected allele pairs with strong LD for structural and interaction analysis: *HLA-A*\**11:01:01:01* with *KIR2DL1*\**0030204, HLA-DPB1*\**05:01:01:01* with *KIR2DL1*\**0030205, CYP2D6*\**1* with *KIR2DP1*\**0020119*, and *CYP2D6*\**1* with *HLA-A*\**01:01:01:01* (Figure 7e-n). Protein structures were predicted using ESM Fold^72^, and binding affinities were evaluated with PRODIGY at 25°C^72,73^. The proteins of all allele pairs demonstrated high binding affinities (*ΔG*, kcal/mol) and low dissociation constants (Kd, M), indicative of strong molecular interactions. The proteins of *CYP2D6*\**1* and *HLA-A*\**01:01:01:01* exhibited the strongest binding (*ΔG* = −231.6 kcal/mol, Kd = 1.4 × 10^−170^), supported by the highest number of intermolecular contacts (3,105) and substantial contributions from charged (28.22%) and apolar residues (44.81%). The proteins of *CYP2D6*\**1* and *KIR2DP1*\**0020119* also displayed strong binding (*ΔG* = −152.8 kcal/mol, Kd = 8.6 × 10^−113^), reflecting extensive structural compatibility. Similarly, the proteins of *HLA-A*\**11:01:01:01* and *KIR2DL1*\**0030204* (*ΔG* = −103.9 kcal/mol, Kd = 6.3 × 10^−77^) and the proteins of *HLA-DPB1*\**05:01:01:01* and *KIR2DL1*\**0030205* (*ΔG* = −56.6 kcal/mol, Kd = 2.9 × 10^−42^) exhibited substantial binding strength, with hundreds to thousands of intermolecular contacts, underscoring the significant compatibility of these molecular interactions. Across all gene pairs, stronger protein interactions, such as those in the *CYP2D6-HLA* and *CYP2D6-KIR* pairs, were associated with higher densities of intermolecular contacts and significant contributions from both charged and apolar residues. The *HLA-A-KIR* and *HLA-DPB1-KIR* pairs, while having fewer contacts, also demonstrated binding affinities that reflect meaningful structural compatibility and stability.

These results suggest that molecular interactions likely contribute to the strong LD between these gene families. The consistently high binding affinities across *CYP2D6-HLA, CYP2D6-KIR, HLA-A-KIR*, and *HLA-DPB1-KIR* proteins underscore their potential role in stabilizing associations within these loci.

### SpecImmune facilitates the discovery of *de novo* mutations

To evaluate SpecImmune’s proficiency in identifying *de novo* mutations (DNMs), we analyzed its typing outcomes in parent-offspring trios from the 1kGP dataset (see Supplementary Table S2). Within the SH006 trio, DNMs were unearthed at the *HLA-DOB* locus in the child. By comparing the child’s inherited haplotypes with those of the parents at each gene locus, discrepancies were flagged as DNMs. To validate these findings, we illustrated the allele sequences of the trio with supporting reads in Figure 8 and Supplementary Figure S14-16. These discrepancies between the child’s haplotype and the parental haplotypes signify recent genomic alterations in the child. Notably, consistent support from the child’s sequencing reads reinforces the credibility of the DNMs pinpointed by SpecImmune. SpecImmune stands out in reconstructing personalized haplotypes and can detect DNMs within immune-related loci.

**Figure 8.**
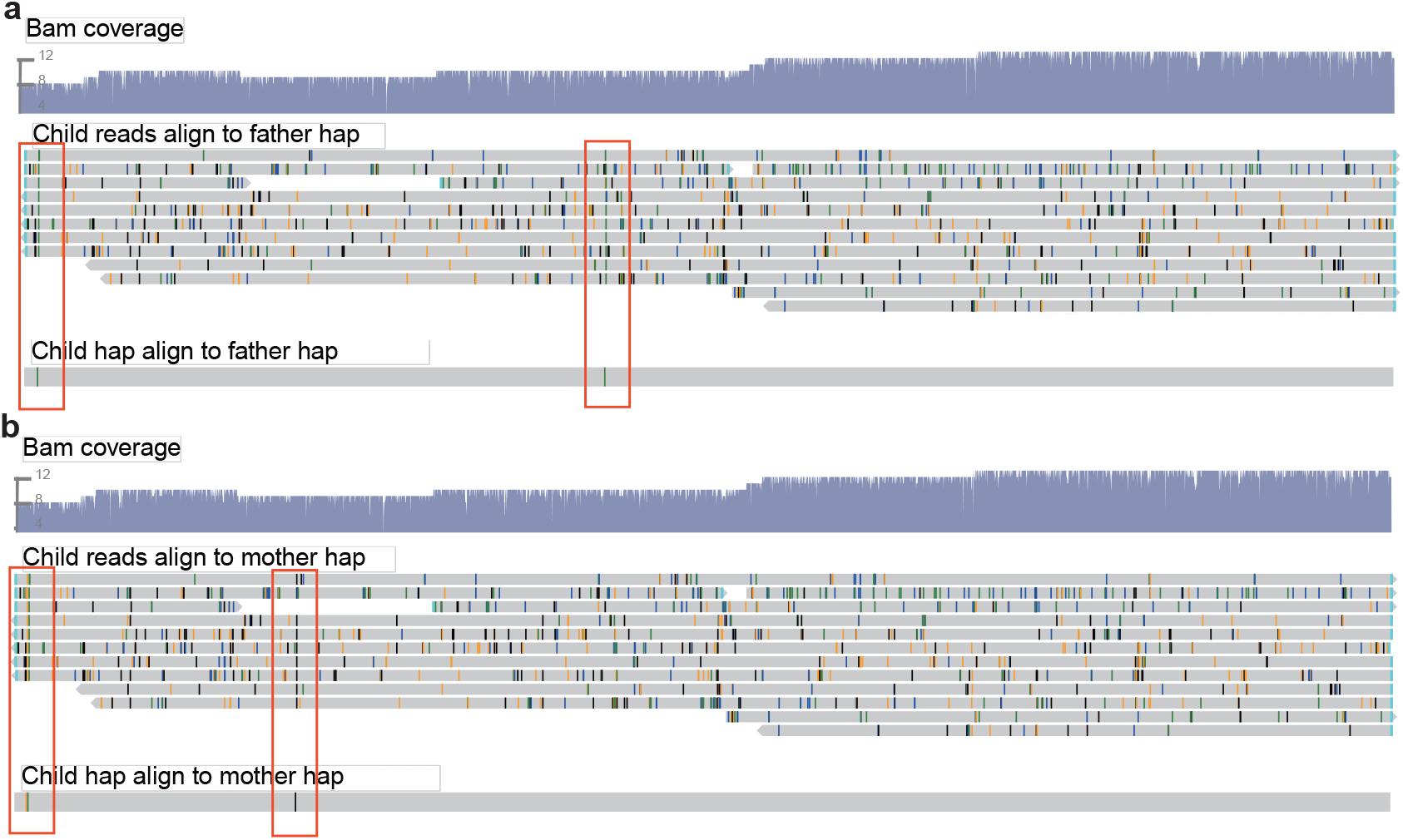
Illustration of DNMs detected by SpecImmune. DNMs are identified at the locus *HLA-DOB* within the family trio SH006. The child is homozygous at this locus. The figure presents the child’s haplotype aligned with the haplotypes of the father (a) and mother (b). Only the inherited haplotype is displayed. In each subplot, the top track illustrates the coverage of the reads alignment, the second track displays the alignment of the child’s reads onto the parent’s haplotype, and the third track showcases the alignment of the child’s haplotype onto the parent’s haplotype. Variants are highlighted in color within the alignment.

### SpecImmune enables clinical-grade personalized allele-based drug recommendations

SpecImmune enables clinical-grade, personalized drug guidance, playing a critical role in advancing precision medicine. By integrating PharmGKB’s clinical guidelines and FDA drug label annotations^74^, it offers allele-based dosing recommendations tailored to individual genetic profiles. Its comprehensive database, encompassing 154,472 allele annotation entries from the HLA and CYP gene families, links alleles to clinically relevant phenotypes such as drug toxicity and efficacy, supported by evidence scores to ensure reliability and applicability^75^.

In precision medicine, accurate genotyping is essential for optimizing drug metabolism and treatment regimens. For sample HG00673, SpecImmune correctly identified the *CYP2D6* genotype as \**1/*\**10*, while pangu misclassified it as \**1/*\**75*. Given the \**10* allele’s decreased function (activity score 0.25) and the \**75* allele’s even lower activity, this error could lead to inappropriate dosing adjustments, undermining treatment safety and efficacy. Similarly, for sample HG01071, SpecImmune accurately identified the *CYP2D6* genotype as \**4/*\**41*, corresponding to an intermediate or poor metabolizer phenotype, while pangu reported it as \**39/*\**39*. Misclassifying \**4/*\**41* (no-function and decreased-function alleles) as \**39/*\**39* (normal-function equivalent) could significantly overestimate metabolic capacity, risking drug toxicity or therapeutic failure for narrow therapeutic index drugs like tricyclic antidepressants or opioids. By reliably identifying genotypes for highly polymorphic genes like *CYP2D6*, SpecImmune enables precise, genotype-guided treatment decisions, directly supporting the goals of precision medicine.

## Methods

### Allele database collection

The allele database serves the purpose of read binning and allele sequence nomenclature within the SpecImmune tool. We obtained the IPD-IMGT/HLA v3.56.0 database^76^ for HLA (https://www.ebi.ac.uk/ipd/imgt/hla/index.html), the IPD-KIR v2.13.0 database^77^ for KIR (https://www.ebi.ac.uk/ipd/kir/), the IMGT/V-QUEST database^78^ (2024-07-10) for IG and TCR (https://www.imgt.org/IMGTindex/V-QUEST.php), and the PharmVar v6.1.2.1 database^79^ for CYP (https://www.pharmvar.org/).

### SpecImmune Algorithm

#### Reads binning

To enhance accuracy and reduce the computational complexity in genotyping, we allocate reads to their respective gene loci, building upon our prior HLA-typing approach, SpecHLA^29^. This methodology assumes that a read exhibits the highest identity to the allele from its originating locus when compared to alleles from other loci. For WGS data, we extract reads from specific regions such as HLA and KIR based on their alignment positions on the hg38 reference genome (Figure 1a-b). Subsequently, these extracted reads are aligned to all alleles in the database. SpecImmune supports BWA MEM^80^ and Minimap2^81^ as alignment methods. During each read alignment, the identity between the read and the mapped allele is computed as

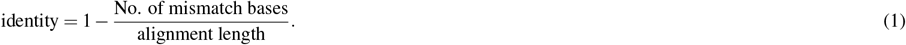

The highest identity among all alleles of a gene locus is designated as the identity for the locus. Subsequently, a read is allocated to the gene locus with the highest identity (Figure 1c). Those with an identity lower than a specific threshold (default set to 85%) were excluded to mitigate noisy reads.

A long read can span multiple gene loci. SpecImmune can assign a read to multiple gene loci to address this scenario. A criterion assigns a read to multiple loci to ensure precision. The distance between every pair of gene loci on the hg38 reference genome is logged. When a read maps to two gene loci, the mapping intervals on the read for the two loci are denoted as [*s*_1_, *e*_1_] and [*s*_2_, *e*_2_]. The distance between the alignments of the two loci on the read (*D*_*reads*_) can be calculated as

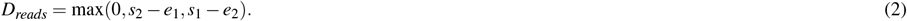

Assuming the distance between the two loci on the hg38 reference is denoted as *D*_ref_, we postulate that the gene distance on hg38 and in an individual genome should align closely. This assumption serves as a constraint for read assignments. If |*D*_*reads*_ − *D*_*re f*_ |< *ϑ* and *D*_*reads*_ > 0, the read is assigned to these two loci. Here, *ϑ* defaults to 2,000. Otherwise, the read is only assigned to the locus with the highest identity.

#### Best-matched alleles selection

A pair of alleles is meticulously chosen from the database for each gene locus to optimize the fit with the reads binned to it. If the allele pair exists in an individual’s genome, the reads will be mapped to it with greater identity and a higher number of matched DNA bases than other allele pairs. We select the best-matched allele pair based on this assumption. We select a condensed candidate set of alleles from the database for each locus to decrease computational complexity. Within each locus, reads are aligned to all locus alleles in the database using Minimap2^81^ with default parameters. The alignment depth of each allele is tallied. Subsequently, only the top 200 alleles with the highest depth are chosen to create the condensed candidate allele set. Then, the reads are realigned to the condensed candidate allele set using Minimap2^81^ with the parameter -p 0.1 -N 100000. This parameter is expressly set to retain all potential alignment records for every read. Three matrices, *M*^*R*×*L*^, *S*^*R*×*L*^, and *D*^*R*×*L*^, are established to house alignment metrics between each read *r* ∈ R and each allele *a* ∈ L. When a read aligns to an allele, *M*^*R*×*L*^, *S*^*R*×*L*^, and *D*^*R*×*L*^ store the count of matched bases, the count of mismatched bases, and the alignment identity, respectively (Figure 1d).

Subsequently, an appropriate allele pair is determined by maximizing the total count of matched bases and the alignment identity across all reads. For every pair of alleles *a*_*i*_ and *a* _*j*_, the aggregate counts of matched bases (*α*_*i j*_), mismatched bases (*β*_*i j*_), and alignment identity (*γ*_*i j*_) from all reads are computed (Figure 1d). This computation assigns each read to the allele with the higher identity. Specifically, if 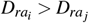, the read *r* is assigned to allele *a*_*i*_; if 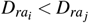, it is assigned to allele *a* _*j*_. In the case of 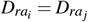, the read *r* is assigned randomly with an equal probability. Let *ξ*_*r*_ = *i or j* store which allele the read *r* is assigned to. Only the alignment between the read and the assigned allele is utilized in determining the values of *α*_*i j*_, *β*_*i j*_, and *γ*_*i j*_ for the given allele pair. The calculation is as follows:

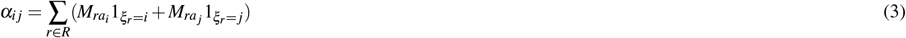

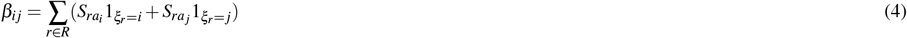

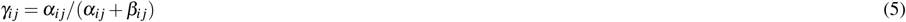

An empirical approach is utilized to determine the optimal allele pair by effectively balancing the maximization of matched bases and alignment identity. *α, β*, and *γ* values are computed for each pair of alleles. The allele pairs are then arranged based on the *α* values. Those pairs with *α* values lower than the highest *α* value by a specified cutoff *τ* form a candidate set. Within this candidate set, the allele pairs are further sorted according to the *γ* values. The highest *γ* value is selected as the best-matched allele pair (Figure 1e).

The beta distribution is employed to determine the heterozygosity or homozygosity of a gene locus initially. For the selected allele pair (*a*_*i*_ and *a* _*j*_), we calculate the depth of each allele (*d*_*i*_ and *d* _*j*_) based on the reads assigned to them. Assuming the lengths of the two alleles are *l*_*i*_ and *l* _*j*_, *d*_*i*_ and *d* _*j*_ are computed as:

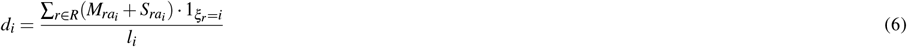

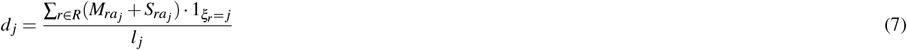

The frequencies of the two alleles are estimated using the beta distribution *B*(*α, β*). The frequency of allele *a*_*i*_ (*κ*_*i*_) is given by:

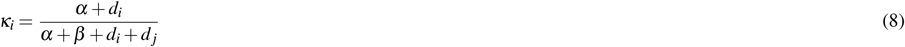

Similarly, the frequency of allele *a* _*j*_ (*κ*_*j*_) is calculated as:

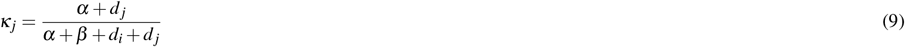

By default, *α* and *β* are both set to 1. Subsequently, the minor allele frequency (*MAF*) is determined as min(*κ*_*i*_, *κ*_*j*_). The locus is considered homozygous if *MAF* < *χ* and heterozygous otherwise, where *χ* defaults to 0.3. The following step utilizes the identified best-matched alleles as personalized reference alleles.

#### Personalized haplotypes reconstruction

We reconstruct personalized haplotypes by leveraging the reads aligned to the best-matched personalized reference alleles to address this. At first, variants are identified based on the reads alignment for both heterozygous and homozygous gene loci. We use Longshot v1.0.0^54^ to call single nucleotide variants (SNVs) and Sniffles v2.2^55^ to detect structural variant (SV) breakpoints. Additionally, SpecImmune supports DeepVariant^82^ as an alternative small variant caller. Since the reads are already phased, we only perform variant calling without further phasing for heterozygous gene loci. Alle-specific reads are aligned separately to the two best-matched personalized reference alleles to identify variants. For homozygous gene loci, as the heterozygous gene loci might be falsely identified as homozygous, these regions may contain heterozygous variants. Therefore, we first perform variant calling, followed by phasing. In this case, locus-specific reads are aligned to the single best-matched personalized reference allele. Subsequently, haplotype phasing of SNVs is conducted using Whatshap v2.3^56^, while joint phasing of both SNVs and SV breakpoints is performed with Longphase v1.7.3^57^.

Next, we reconstruct personalized linear haplotype sequences based on the identified variants. We first directly generate consensus sequences based on phased SNVs on the personalized reference allele for each haplotype. After that, since the connections between phased SV breakpoints on the same haplotype could be non-linear (Figure 1f3), a *haplotype reconstruction* algorithm is applied to obtain entirely linear haplotypes. We segment the personalized reference allele and link the segments to reconstruct the individual haplotype (Figure 1g). We segment the consensus sequences based on the SV breakpoints for each haplotype and then estimate the copy number of each segment. The resulting segments are denoted as *S* = {*s*_1_, *s*_2_, …, *s*_*n*_}, where *n* represents the number of segments.

SpecImmune estimates the copy numbers of genomic segments, denoted as {*c*_1_, *c*_2_, …, *c*_*n*_}, using a two-step approach. First, the mean read alignment depth across all *n* segments is calculated as:

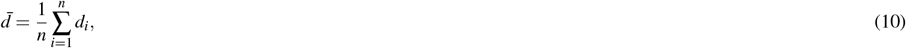

where *d*_*i*_ is the read alignment depth of segment *i*. Second, the copy number *c*_*i*_ of segment *i* is estimated as:

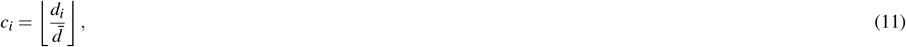

where ⌊·⌋ denotes the floor function. The weights of the edges are set to the read counts derived from spanning reads, which quantify the strength of adjacency between segments. SpecImmune uses Minimap2^81^ to align and identify all reads spanning two segments. After constructing the graph, we adopt the *haplotype reconstruction*^58,59^ algorithm to reconstruct the haplotype sequence^58,59^.

SpecImmune iteratively performs haplotype sequence reconstruction; we realign the allele-specific reads to the haplotype sequences obtained from the *haplotype reconstruction*^58,59^. Following this, we perform variant calling, phasing, segmentation, and *haplotype reconstruction*^58,59^. This entire process is iteratively repeated until no variant is identified. Then, the personalized diploid haplotype sequences are reconstructed for each locus. To mitigate noise resulting from low sequencing depth, a window-sliding method is applied to mask low-depth regions, inspired by previous work^29^. This method involves sliding a 20 bp window along the gene locus, masking any window with a mean depth below a specified threshold value (defaulting to five) with the character ‘N’ (Supplementary Algorithm 1).

#### Nomenclature

SpecImmune assigns an official designation to each personalized haplotype sequence, corresponding to the best-matched alleles retrieved from the allele database (Figure 1e). This assignment thoroughly compares the reconstructed sequence and all alleles cataloged in the database, utilizing Blastn v2.15.0^83^. An empirical approach is utilized to select the best-matched alleles. For each allele, Blastn reports the number of mismatches (*m*), the length of gaps (*g*), and the mapped length (*L*). The identity (*ι*) is calculated as

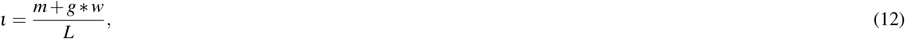

where *w* is a hyperparameter default by 0.4. This parameter is utilized to modulate the significance of gaps in identity calculation. Given that the predominant errors in long reads are insertion/deletion errors^54^, we employ this hyperparameter to adjust the weighting of gaps, thereby mitigating the influence of insertion/deletion errors.

We select the best-matched alleles based on the mapped length and identity to prevent cases where alleles exhibit short local alignments but high identity. The process involves creating a candidate set where alleles with a mapped length lower than the highest mapped length value by a specific cutoff (defaulting to 0.02) are included. This candidate set arranges alleles based on their identity (*ι*) values. Alleles with an *ι* lower than the highest *ι* value by a specific cutoff (defaulting to 4e-4) are designated as the best-matched alleles. The nomenclature schemes for HLA, KIR, and CYP alleles are depicted in Supplementary Figure S1.

#### Report visualization

To depict DNMs and demonstrate the confidence level of the reported alleles, we visualize the typing alleles and supporting reads within SpecImmune’s output report. We have developed a visualization module based on the GenomeView engine^84^. By default, the visualization report comprises six tracks: sample information, results track, coverage track, variant track, alignment track, and candidate alleles track (Figure1h). The module incorporates various features to improve the visualization of noisy long-read sequencing data, including from PacBio and ONT platforms. Notably, it consists of a quick-consensus mode that masks potential sequencing errors while highlighting probable variants. The module supports output in high-quality formats such as SVG, PDF, and PNG, facilitating the automated generation of visualizations within analysis pipelines.

#### IG/TCR typing

Given the relatively short nature of IG and TCR alleles, binning reads to the gene locus poses a challenge. We have devised a specialized pipeline for IG/TCR typing. Initially, all IG/TCR reads are aligned to the no-alt hg38 reference using Minimap2^81^, which contains only the main chromosomes. Subsequently, SNVs are called using Longshot v1.0.0^54^, and SVs are called using pbsv (https://github.com/PacificBiosciences/pbsv). The variant files of SNVs and SVs are merged using bcftools (https://samtools.github.io/bcftools/bcftools.html). Following this, all variants undergo phasing using Whatshap 2.3 ^56^. Based on the phased variants, two haplotype sequences are then generated for each gene locus. Regions with low sequencing depth are masked utilizing the method above. We deduce the nomenclature for each haplotype sequence employing the specified method. Furthermore, we provide the phase set at the locus for loci with heterogeneous variants. This conveys the phase information between different gene loci.

#### CYP typing

Copy number variations (CNVs) and gene fusions are commonly observed at the *CYP2D6* locus, presenting a challenge for our core typing method. To tackle this challenge, we have devised a strategy that integrates the results of pangu with our core method for *CYP2D6* typing (https://github.com/PacificBiosciences/pangu). We have adapted the pangu approach with minor adjustments to effectively type alleles that exhibit CNVs and gene fusions. Both our core typing method and pangu are executed independently. To utilize pangu, reads are aligned to the non-alt hg38 reference. To enhance typing accuracy and mitigate potential errors and ambiguities in the alignments, we refine the alignments as follows. Initially, alignments with an identity below 0.85 are excluded from further analysis. Additionally, in scenarios where alleles may result from a fusion of *CYP2D6* and *CYP2D7*, reads are assigned to the gene with the higher identity. Given pangu’s optimization for PacBio HiFi data, we have enriched it with additional allele markers from various data protocols to cater to diverse data sources. If pangu’s results suggest the presence of CNVs or gene fusions, we consider pangu’s outputs as the final result. Conversely, if the locus represents a standard diploid gene without such variations, we rely on the results generated by our core typing method. The typing of other CYP loci is exclusively conducted utilizing our core algorithm.

We employ the previous Stargazer^40^ method to forecast the phenotype using the star alleles as a foundation. *CYP2D6* haplotypes are translated into the standard unit of enzyme activity, referred to as an activity score. These activity scores play a crucial role in predicting the four metabolizer classes, categorized as follows: poor metabolizers with a score of 0; intermediate metabolizers with a score of 0.5; normal metabolizers ranging from 1 to 2; and ultrarapid metabolizers with a score exceeding 2.

### RNA-based HLA typing

For RNA sequencing data, we align locus-specific reads to the hg38 reference genome and perform variant calling using Longshot v1.0.0. We then phase the variants using WhatsHap v2.3^85^. Subsequently, allele-specific reads are generated by leveraging WhatsHap haplotag assignments. These allele-specific reads are assembled in a reference-guided manner using StringTie version 2.2.3^56^. Finally, we generate diploid haplotype sequences by masking intronic regions with ‘N’s.

### Benchmark of SpecImmune

#### Benchmark data

To validate SpecImmune, we gathered a 1kGP dataset^51^, two HPRC datasets, and two HGSVC datasets (Table 2). The 1kGP dataset comprises 1,019 ONT WGS samples. The reference HLA types of 1kGP individuals were acquired from https://ftp.1000genomes.ebi.ac.uk/vol1/ftp/data_collections/HLA_types/. Furthermore, we obtained a HiFi WGS and an ONT WGS dataset from the HPRC project^52^. Additionally, we collected a HiFi WGS dataset and a PacBio CLR WGS dataset from the HGSVC project^53^. Phased assemblies are accessible for samples from the HPRC and HGSVC projects. Sequencing reads of HLA, KIR, IG, TCR, and CYP loci were extracted by aligning the WGS data to the hg38 reference. For CYP typing validation, we also gathered 10 PacBio HiFi amplicon datasets with reference *CYP2D6* alleles from prior research^42^. The reference *CYP2D6* alleles of HPRC datasets were obtained from previous research, inferred from matched short-read data (https://pacbio.cn/wp-content/uploads/poster_harting.pdf). The reference *CYP2D6* alleles in the 1kGP dataset were sourced from the GeT-RM project^62^. Additionally, we collected RNA sequencing data for HG002 LCL and iPSCs, and HG004 and HG005 LCLs, including PacBio Iso-seq, MAS-seq, and ONT direct RNA, from the GIAB project^60^. The detailed command parameters employed for SpecHLA, HLA*LA, and SpecImmune are available in Supplementary Table S1.

Additionally, we employed simulated data to assess the performance of SpecImmune. We randomly chose a pair of alleles for each sample for each locus. Subsequently, we generated long reads for each selected allele sequence using PBSIM3^61^ with the configuration `-hmm_model P4C2.model`. Finally, the simulated reads were consolidated into a single `fastq` file for each sample.

#### Allele annotation on phased genomes

To evaluate SpecImmune, we deduce the reference alleles from the phased assemblies of the HPRC and HGSVC samples. For identifying alleles of the intricate genes on a phased reference genome, we align the assembled genome to the allele database of HLA, KIR, IG, TCR, and CYP loci. We employed BWA-MEM for the alignment process to achieve precise base-level alignments, as referenced in prior work^23^. Subsequently, reference alleles were chosen based on these alignments.

Reference alleles were meticulously selected to optimize alignment length and identity with the phased assembly. For each gene, we compiled the alignment ratio (*η*), alignment length, and alignment identity of all alleles within the gene locus. The alignment ratio, calculated as the alignment length divided by the allele’s length, excluded alleles with *η* < 0.95. Next, a candidate set was created, encompassing alleles with an alignment length lower than the highest value by a specified cutoff. Within this set, alleles were organized based on their identity values, retaining those below the highest identity value by a designated cutoff. These curated alleles constitute the reference set used to evaluate the accuracy of SpecImmune.

#### Benchmark Metrics

The typing accuracy is defined as the ratio of correctly inferred alleles to all inferred alleles. To account for ambiguities in the ground truth and the inferred gene types when counting correctly inferred alleles, we employ a compatible function *F* to evaluate typing accuracy, as outlined in prior work^29^. The function *F*(*R, I*) yields a value of 1 only if the intersection between sets *R* and *I* is not empty. Considering a locus, where the reference gene type tuples are denoted as *R*_1_ and *R*_2_, and the inferred gene type tuples are represented by *I*_1_ and *I*_2_, the calculation for the number of correctly inferred alleles involves determining

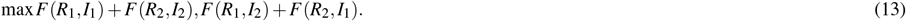

We employ the identical allele database version throughout the evaluation to maintain nomenclature consistency. Given that the HLA typing method HLA*LA prohibits database version alterations, we align the results from HLA*LA to our IMGT/HLA database version. Furthermore, we harmonize the reference HLA alleles reported by the 1kGP to our IMGT/HLA database version to ensure uniformity.

### Gene allele diversity quantification

The Shannon Diversity index is used to quantify the allele diversity at each gene locus. Denoted as *H*, the Shannon Diversity index can be calculated as:

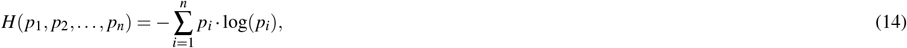

where *p*_1_, *p*_2_, …, *p*_*n*_ are the proportions of different alleles at the gene locus.

### Categorization of alleles by frequency

Drawing from prior research^86^, we delineated alleles into distinct categories based on their occurrence frequencies: common alleles (> 5%), low-frequency alleles (between 0.5% and 5%), and rare alleles (< 0.5%). At each locus, the allele frequency is determined by dividing the number of a specific category by the total number of alleles identified within the population.

### Quantify allele frequency difference between populations

Utilizing SpecImmune, we derive a set of alleles inferred from all 1kGP samples. For each allele, we calculate its frequency across all 26 populations. Subsequently, we quantify the disparity in allele frequencies between every pair of populations. Considering two populations, let *p*_*i*_ represent the *i*-th allele in the first population, and *q*_*i*_ represent the *i*-th allele in the second population. Assuming there are a total of *n* alleles, we have 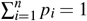 and 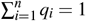. The distinction in allele frequencies between the two populations is assessed using the Jensen-Shannon distance (JSD), which is computed as:

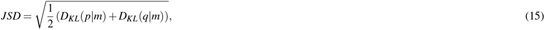

where *m* denotes the pointwise mean of *p* and *q*, and *D*_*KL*_ signifies the Kullback-Leibler divergence. This calculation uses the `distance.jensenshannon` function provided in the Python `Scipy` package.

### LD analysis

For each pair of loci, we computed the linkage disequilibrium based on the alleles inferred by SpecImmune using the R package `pould` v1.0.1^65,66^. The typed alleles are in the highest resolution, specified at a four-field level for HLA alleles and a three-field level for KIR alleles. The alleles across distinct loci are not phased. To estimate LD for unphased loci, we utilized the function `cALD(data, inPhase=FALSE)`. The metric *Wn* quantifies the degree of LD. It ranges from 0 to 1, where *Wn* = 0 signifies complete independence, while *Wn* = 1 indicates complete LD. Also, the pair of conditional asymmetric LD (ALD) measures, *W*_*b/a*_ and *W*_*a/b*_, are used to complement the *Wn* measure due to there are different numbers of alleles at the two loci *a* and *b*^65^. Furthermore, we define the minimal ALD value *minALD*, which is calculated as *minALD* = *min*(*W*_*b/a*_,*W*_*a/b*_).

### Differential IG/TCR loci of heterozygous variants

SpecImmune infers the count of heterozygous variants at each IG/TCR locus. Subsequently, we identify the differential IG/TCR loci of heterozygous variants, characterized by significantly divergent counts of heterozygous variants between populations. A vector is allocated to each locus to document the number of heterozygous variants in every sample in a group. We assess the two vectors for each locus using a t-test when comparing two groups. t-test is achieved by the Python `scipy.stats.ttest_ind` function. The resulting p-values are adjusted utilizing the Bonferroni method. This is implemented by the Python `statsmodels.stats.multitest.multipletests` function. Loci with adjusted p-values below 0.01 are designated as significant differential loci.

### *De novo* mutations discovery

To identify *De novo* mutations (DNMs), we explored the mutations that violated Mendelian inheritance in the family trios. We aligned the haplotype sequences obtained using SpecImmune to the human reference genome (hg38). Variant calling, including SNVs and small insertions and deletions (indels), was performed using samtools mpileup and bcftools call. Mutations supported by at least five reads in WGS data from the 1000 Genomes Project https://www.internationalgenome.org/data-portal/sample were retained. Subsequently, we identified mutation sites within each family that violated Mendelian inheritance^87–89^. The total number of such sites is represented as *N*_DNMs_, and the total number of genomic sites analyzed is denoted as *N*_total_. The DNMs rate is defined as:

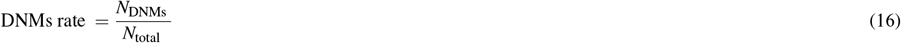

## Discussion

In HLA genotyping from sequencing data, the conventional method of selecting the best-matched allele pair from the database based on sequencing reads alignment is a well-established approach known for its reliability. For instance, OptiType^18^ maximizes read alignment to chosen alleles using integer linear programming, while PolySolver^19^ integrates aligned read base qualities and observed insert sizes through a Bayesian model to select alleles. HLA-VBSeq^21^ optimizes read alignment and quantities to alleles using variational Bayesian inference. However, existing methods following this strategy are primarily tailored for short reads sequencing data. Long reads present distinct challenges compared to short reads due to their extended length, higher error rates, and predominantly single-end. The highly variable length of long reads presents a challenge for existing methods. Despite benefiting from their length, long reads can map to nearly all alleles of a specific gene locus, making it impractical to rely solely on maximizing the number of reads mapped to inferred alleles. Our approach proposes a novel method to maximize the alignment identity between reads and alleles. Given that a read is typically derived from a specific allele, the alignment identity between the read and its source allele should surpass that of other alleles. However, a brief partial alignment between the read and a non-source allele may demonstrate a higher level of identity than a full-length alignment between the read and the source allele. To address this challenge, we introduce the concept of matched base numbers. Our strategy involves selecting an allele pair that maximizes alignment identity and matched base numbers for all reads. Benchmark tests have demonstrated the effectiveness and reliability of this strategy for choosing the best-matched allele pair from long reads data.

SpecImmune employs a read-binning strategy for genotyping that involves three key steps. Initially, it extracts reads from a specific region of interest, bins them into respective gene loci, and finally assigns them to specific alleles. By maximizing alignment identity and matching base numbers between all binned reads and alleles, SpecImmune accurately infers gene types. Through continual read binning, this method effectively reduces computational complexity. Extensive benchmark experiments have consistently verified the reliability of this approach for genotyping from long-read data, focusing on HLA, KIR, IG, TCR, and CYP genes. Notably, this approach is versatile and can be applied to typing other genes.

In addition to the HLA genes, several other immune-related genes within the HLA region contribute significantly to immune regulation. For example, *HFE* encodes a membrane protein similar in structure to HLA class I molecules and plays a key role in regulating iron balance^90^. *MICA* and *MICB* are stress-induced antigens that serve as ligands for the activating receptor NKG2D, which is expressed in NK and T cells^91^. *TAP1* and *TAP2*, members of the ATP-binding cassette transporter family, are critical for transporting peptide antigens into the endoplasmic reticulum, where they are loaded onto HLA class I molecules for presentation to CD8+ T cells^92^.

SpecImmune can conduct gene typing swiftly, often completing the process within a few hours. By harnessing long-read real-time sequencing, SpecImmune can acquire real-time data, resulting in prompt immune-related gene typing outcomes. This real-time functionality proves particularly beneficial in urgent clinical scenarios such as emergency organ transplant matching. Furthermore, SpecImmune offers reconstructed allele sequences, and visual representations of aligned reads supporting these sequences, facilitating users in evaluating the accuracy of the typing outcomes. In addition to handling HLA class I and II genes, SpecImmune extends its capabilities to include HLA class I pseudogenes and non-HLA genes within the MHC region. SpecImmune can also type KIR, IG, TCR, and CYP genes. This breadth of coverage renders SpecImmune valuable for clinical applications and immune-related research endeavors.

Our study faces two primary limitations. Firstly, our study predominantly focuses on typing individual gene loci, lacking information on phasing across different loci. We only inferred the haplotype among genes for IG/TCR typing. Long reads offer valuable long-range phasing details, presenting an opportunity to reconstruct haplotypes spanning multiple gene loci. Such haplotypes hold significance for medical research and evolutionary studies. Secondly, our core methodology does not account for gene fusion events because we concentrate independently on each gene locus. Complex genes like *CYP2D6* often feature gene fusion events. To address these variants within *CYP2D6*, we incorporate the third-party tool pangu. The pangu employs an empirical approach that utilizes specific SNPs or SV breakpoints to infer allele types. Dealing with novel alleles necessitates utilizing a prior knowledge-independent method to manage gene fusion events effectively.

## Data availability

The 1KG-ONT panel was retrieved from https://ftp.1000genomes.ebi.ac.uk/vol1/ftp/data_collections/1KG_ONT_VIENNA/. Sequencing data for the HPRC and HGSVC projects can be accessed at https://humanpangenome.org/ and https://www.internationalgenome.org/data-portal/data-collection/hgsvc3, respectively. Corresponding assemblies are available at https://github.com/human-pangenomics/HPP_Year1_Assemblies for HPRC and https://www.hgsvc.org/resources for HGSVC. GIAB RNA sequencing data can be found at https://ftp-trace.ncbi.nlm.nih.gov/ReferenceSamples/giab/data_RNAseq/.

## Code availability

The software package SpecImmune is freely available at https://github.com/deepomicslab/SpecImmune. The scripts required to reproduce the results are also available in this repository.

## Funding

This work was supported by the Shenzhen Science and Technology Program (20220814183301001).

### Acknowledgements

We express our gratitude for the generous support provided by the Shenzhen Science and Technology Program (Grant No. 20220814183301001). We acknowledge the Institute of Molecular Pathology (Vienna, Austria) for publishing the 1KG-ONT panel. We acknowledge the use of *BioRender.com* for assistance in creating the Figure 1 used in the visualization of the workflow. Figures were generated using *BioRender* under a publication license. © Wang, X. (2025). Retrieved from https://BioRender.com/y24b592. We extend our gratitude to the contributors of the HPRC and HGSVC projects. Special thanks to Dr. Yiqi Jiang for her invaluable advice on figure design. Additionally, we acknowledge Lijia Che for her contribution to visualizing the LD graph.

## Author contributions statement

S.C.L. designed and supervised the study. S.W. and X.D.W. developed the software and wrote the manuscript. M.Y.W. was involved in the software development. Q.Z. was involved in the drug recommendation procedure development and software test. All authors contributed to the manuscript revisions.

## References

1. James Robinson, Jason A Halliwell, James D Hayhurst, Paul Flicek, Peter Parham, and Steven GE Marsh. The ipd and imgt/hla database: allele variant databases. Nucleic Acids Res., 43(D1):D423–D431, 2015.

2. Carlos Vilches and Peter Parham. Kir: Diverse, rapidly evolving receptors of innate and adaptive immunity. Annual Review of Immunology, 20(Volume 20, 2002):217–251, 2002.

3. Sergei Radaev and Peter D. Sun. Structure and function of natural killer cell surface receptors*. Annual Review of Biophysics, 32(Volume 32, 2003):93–114, 2003.

4. Peter Parham. Immunogenetics of killer cell immunoglobulin-like receptors. Molecular Immunology, 42(4):459–462, 2005. Natural Killer cells from ‘disturbing’ background to central players of immune responses.

5. David G. Schatz and Patrick C. Swanson. V(d)j recombination: Mechanisms of initiation. Annual Review of Genetics, 45(Volume 45, 2011):167–202, 2011.

6. Mark M Davis. T-cell antigen receptor genes and t-cell recognition. Nature, 334:395–402, 1988.

7. F. Peter Guengerich. Cytochrome p450 research and the journal of biological chemistry. Journal of Biological Chemistry, 294(5):1671–1680, 2019.

8. F. Peter Guengerich. Role of cytochrome p450 enzymes in drug-drug interactions. volume 43 of Advances in Pharmacology, pages 7–35. Academic Press, 1997.

9. Patrick J. Medina and Val R. Adams. Pd-1 pathway inhibitors: Immuno-oncology agents for restoring antitumor immune responses. Pharmacotherapy: The Journal of Human Pharmacology and Drug Therapy, 36(3):317–334, 2016.

10. Elisavet Stavropoulou, Gratiela G Pircalabioru, and Eugenia Bezirtzoglou. The role of cytochromes p450 in infection. Frontiers in immunology, 9:89, 2018.

11. Bryan Briney, Anne Inderbitzin, Collin Joyce, and Dennis R. Burton. Commonality despite exceptional diversity in the baseline human antibody repertoire. Nature, 566:393–397, 2019.

12. Brahma V. Kumar, Thomas J. Connors, and Donna L. Farber. Human t cell development, localization, and function throughout life. Immunity, 37(2):202–213, 2018.

13. Ashley Moffett and Francesco Colucci. Co-evolution of nk receptors and hla ligands in humans is driven by reproduction. Immunological Reviews, 267(1):283–297, 2015.

14. Peter Parham and Ashley Moffett. Variable nk cell receptors and their mhc class i ligands in immunity, reproduction and human evolution. Nature Reviews Immunology, 13:133–144, 2013.

15. Ronald M. Sobecks, Tao Wang, Medhat Askar, Meighan M. Gallagher, Michael Haagenson, Stephen Spellman, Marcelo Fernandez-Vina, Karl-Johan Malmberg, Carlheinz Müller, Minoo Battiwalla, James Gajewski, Michael R. Verneris, Olle Ringdén, Susana Marino, Stella Davies, Jason Dehn, Martin Bornhäuser, Yoshihiro Inamoto, Ann Woolfrey, Peter Shaw, Marilyn Pollack, Daniel Weisdorf, Jeffrey Milller, Carolyn Hurley, Stephanie J. Lee, and Katharine Hsu. Impact of kir and hla genotypes on outcomes after reduced-intensity conditioning hematopoietic cell transplantation. Biology of Blood and Marrow Transplantation, 21(9):1589–1596, 2015.

16. Katsumi Maenaka, Takeo Juji, Takahiro Nakayama, Jessica R. Wyer, George F. Gao, Taeko Maenaka, Nathan R. Zaccai, Akiko Kikuchi, Toshio Yabe, Katsushi Tokunaga, Kenji Tadokoro, David I. Stuart, E. Yvonne Jones, and P. Anton van der Merwe. Killer cell immunoglobulin receptors and t cell receptors bind peptide-major histocompatibility complex class i with distinct thermodynamic and kinetic properties*. Journal of Biological Chemistry, 274(40):28329–28334, 1999.

17. Y Zhou, M Ingelman-Sundberg, and VM Lauschke. Worldwide distribution of cytochrome p450 alleles: A meta-analysis of population-scale sequencing projects. Clinical Pharmacology & Therapeutics, 102(4):688–700, 2017.

18. András Szolek, Benjamin Schubert, Christopher Mohr, Marc Sturm, Magdalena Feldhahn, and Oliver Kohlbacher. Optitype: precision hla typing from next-generation sequencing data. Bioinformatics, 30(23):3310–3316, 2014.

19. Sachet A Shukla, Michael S Rooney, Mohini Rajasagi, Grace Tiao, Philip M Dixon, Michael S Lawrence, Jonathan Stevens, William J Lane, Jamie L Dellagatta, Scott Steelman, et al. Comprehensive analysis of cancer-associated somatic mutations in class i hla genes. Nat. Biotechnol., 33(11):1152, 2015.

20. Yu Bai, Min Ni, Blerta Cooper, Yi Wei, and Wen Fury. Inference of high resolution hla types using genome-wide rna or dna sequencing reads. BMC genomics, 15:1–16, 2014.

21. Naoki Nariai, Kaname Kojima, Sakae Saito, Takahiro Mimori, Yukuto Sato, Yosuke Kawai, Yumi Yamaguchi-Kabata, Jun Yasuda, and Masao Nagasaki. Hla-vbseq: accurate hla typing at full resolution from whole-genome sequencing data. BMC genomics, 16(2):S7, 2015.

22. Shuji Kawaguchi, Koichiro Higasa, Masakazu Shimizu, Ryo Yamada, and Fumihiko Matsuda. Hla-hd: An accurate hla typing algorithm for next-generation sequencing data. Hum. Mutat., 38(7):788–797, 2017.

23. Li Song, Gali Bai, X Shirley Liu, Bo Li, and Heng Li. Efficient and accurate kir and hla genotyping with massively parallel sequencing data. Genome Research, 33(6):923–931, 2023.

24. Alexander T Dilthey, Pierre-Antoine Gourraud, Alexander J Mentzer, Nezih Cereb, Zamin Iqbal, and Gil McVean. High-accuracy hla type inference from whole-genome sequencing data using population reference graphs. PLoS Comput. Biol., 12(10):e1005151, 2016.

25. Alexander T Dilthey, Alexander J Mentzer, Raphael Carapito, Clare Cutland, Nezih Cereb, Shabir A Madhi, Arang Rhie, Sergey Koren, Seiamak Bahram, Gil McVean, et al. Hla* la—hla typing from linearly projected graph alignments. Bioinformatics, 35(21):4394–4396, 2019.

26. Rose Orenbuch, Ioan Filip, Devon Comito, Jeffrey Shaman, Itsik Pe’er, and Raul Rabadan. arcashla: high-resolution hla typing from rnaseq. Bioinformatics, 36(1):33–40, 2020.

27. Heewook Lee and Carl Kingsford. Kourami: graph-guided assembly for novel human leukocyte antigen allele discovery. Genome biology, 19:1–16, 2018.

28. Daehwan Kim, Joseph M Paggi, Chanhee Park, Christopher Bennett, and Steven L Salzberg. Graph-based genome alignment and genotyping with hisat2 and hisat-genotype. Nat. Biotechnol., 37(8):907–915, 2019.

29. Shuai Wang, Mengyao Wang, Lingxi Chen, Guangze Pan, Yanfei Wang, and Shuai Cheng Li. Spechla enables full-resolution hla typing from sequencing data. Cell Reports Methods, 3(9), 2023.

30. Damjan Vukcevic, James A Traherne, Sigrid Næss, Eva Ellinghaus, Yoichiro Kamatani, Alexander Dilthey, Mark Lathrop, Tom H Karlsen, Andre Franke, Miriam Moffatt, et al. Imputation of kir types from snp variation data. The American Journal of Human Genetics, 97(4):593–607, 2015.

31. David Roe and Rui Kuang. Accurate and efficient kir gene and haplotype inference from genome sequencing reads with novel k-mer signatures. Frontiers in immunology, 11:583013, 2020.

32. Wesley M Marin, Ravi Dandekar, Danillo G Augusto, Tasneem Yusufali, Bianca Heyn, Jan Hofmann, Vinzenz Lange, Jürgen Sauter, Paul J Norman, and Jill A Hollenbach. High-throughput interpretation of killer-cell immunoglobulin-like receptor short-read sequencing data with ping. PLoS computational biology, 17(8):e1008904, 2021.

33. Dmitriy A Bolotin, Stanislav Poslavsky, Alexey N Davydov, Felix E Frenkel, Lorenzo Fanchi, Olga I Zolotareva, Saskia Hemmers, Ekaterina V Putintseva, Anna S Obraztsova, Mikhail Shugay, et al. Antigen receptor repertoire profiling from rna-seq data. Nature biotechnology, 35(10):908–911, 2017.

34. Stefan Canzar, Karlynn E Neu, Qingming Tang, Patrick C Wilson, and Aly A Khan. Basic: Bcr assembly from single cells. Bioinformatics, 33(3):425–427, 2017.

35. Ferran Nadeu, Rut Mas-de Les-Valls, Alba Navarro, Romina Royo, Silvia Martín, Neus Villamor, Helena Suárez-Cisneros, Rosó Mares, Junyan Lu, Anna Enjuanes, et al. Igcaller for reconstructing immunoglobulin gene rearrangements and oncogenic translocations from whole-genome sequencing in lymphoid neoplasms. Nature communications, 11(1):3390, 2020.

36. Michael KB Ford, Ananth Hari, Oscar Rodriguez, Junyan Xu, Justin Lack, Cihan Oguz, Yu Zhang, Andrew J Oler, Ottavia M Delmonte, Sarah E Weber, et al. Immunotyper-sr: A computational approach for genotyping immunoglobulin heavy chain variable genes using short-read data. Cell systems, 13(10):808–816, 2022.

37. Ibrahim Numanagić, Salem Malikić, Victoria M Pratt, Todd C Skaar, David A Flockhart, and S Cenk Sahinalp. Cypiripi: exact genotyping of cyp2d6 using high-throughput sequencing data. Bioinformatics, 31(12):i27–i34, 2015.

38. Greyson P Twist, Andrea Gaedigk, Neil A Miller, Emily G Farrow, Laurel K Willig, Darrell L Dinwiddie, Josh E Petrikin, Sarah E Soden, Suzanne Herd, Margaret Gibson, et al. Constellation: a tool for rapid, automated phenotype assignment of a highly polymorphic pharmacogene, cyp2d6, from whole-genome sequences. NPJ genomic medicine, 1(1):1–10, 2016.

39. Ibrahim Numanagić, Salem Malikić, Michael Ford, Xiang Qin, Lorraine Toji, Milan Radovich, Todd C Skaar, Victoria M Pratt, Bonnie Berger, Steve Scherer, et al. Allelic decomposition and exact genotyping of highly polymorphic and structurally variant genes. Nature communications, 9(1):828, 2018.

40. Seung-been Lee, Marsha M Wheeler, Karynne Patterson, Sean McGee, Rachel Dalton, Erica L Woodahl, Andrea Gaedigk, Kenneth E Thummel, and Deborah A Nickerson. Stargazer: a software tool for calling star alleles from next-generation sequencing data using cyp2d6 as a model. Genetics in medicine, 21(2):361–372, 2019.

41. Xiao Chen, Fei Shen, Nina Gonzaludo, Alka Malhotra, Cande Rogert, Ryan J Taft, David R Bentley, and Michael A Eberle. Cyrius: accurate cyp2d6 genotyping using whole-genome sequencing data. The pharmacogenomics journal, 21(2):251–261, 2021.

42. Sarah Charnaud, Jacob E Munro, Lucie Semenec, Ramin Mazhari, Jessica Brewster, Caitlin Bourke, Shazia Ruybal-Pesántez, Robert James, Dulcie Lautu-Gumal, Harin Karunajeewa, et al. Pacbio long-read amplicon sequencing enables scalable high-resolution population allele typing of the complex cyp2d6 locus. Communications Biology, 5(1):168, 2022.

43. Jesse Bruijnesteijn. Hla/mhc and kir characterization in humans and non-human primates using oxford nanopore technolo-gies and pacific biosciences sequencing platforms. HLA, 101(3):205–221, 2023.

44. Ting Hon, Kristin Mars, Greg Young, Yu-Chih Tsai, Joseph W Karalius, Jane M Landolin, Nicholas Maurer, David Kudrna, Michael A Hardigan, Cynthia C Steiner, et al. Highly accurate long-read hifi sequencing data for five complex genomes. Scientific data, 7(1):399, 2020.

45. Joanne D Stockton, Thomas Nieto, Elizabeth Wroe, Anthony Poles, Nicholas Inston, David Briggs, and Andrew D Beggs. Rapid, highly accurate and cost-effective open-source simultaneous complete hla typing and phasing of class i and ii alleles using nanopore sequencing. HLA, 96(2):163–178, 2020.

46. Neema P Mayor, James Robinson, Alasdair JM McWhinnie, Swati Ranade, Kevin Eng, William Midwinter, Will P Bultitude, Chen-Shan Chin, Brett Bowman, Patrick Marks, et al. Hla typing for the next generation. PloS one, 10(5):e0127153, 2015.

47. Jesse Bruijnesteijn, Marit Van der Wiel, Natasja G De Groot, and Ronald E Bontrop. Rapid characterization of complex killer cell immunoglobulin-like receptor (kir) regions using cas9 enrichment and nanopore sequencing. Frontiers in Immunology, 12:722181, 2021.

48. Oscar L Rodriguez, William S Gibson, Tom Parks, Matthew Emery, James Powell, Maya Strahl, Gintaras Deikus, Kathryn Auckland, Evan E Eichler, Wayne A Marasco, et al. A novel framework for characterizing genomic haplotype diversity in the human immunoglobulin heavy chain locus. Frontiers in immunology, 11:2136, 2020.

49. Oscar L Rodriguez, Catherine A Silver, Kaitlyn Shields, Melissa L Smith, and Corey T Watson. Targeted long-read sequencing facilitates phased diploid assembly and genotyping of the human t cell receptor alpha, delta, and beta loci. Cell Genomics, 2(12), 2022.

50. Jia-Yuan Zhang, Hannah Roberts, David SC Flores, Antony J Cutler, Andrew C Brown, Justin P Whalley, Olga Mielczarek, David Buck, Helen Lockstone, Barbara Xella, et al. Using de novo assembly to identify structural variation of eight complex immune system gene regions. PLoS computational biology, 17(8):e1009254, 2021.

51. Jonas A Gustafson, Sophia B Gibson, Nikhita Damaraju, Miranda PG Zalusky, Kendra Hoekzema, David Twesigomwe, Lei Yang, Anthony A Snead, Phillip A Richmond, Wouter De Coster, et al. High-coverage nanopore sequencing of samples from the 1000 genomes project to build a comprehensive catalog of human genetic variation. Genome Research, 34(11):2061–2073, 2024.

52. David Porubsky, Mitchell R Vollger, William T Harvey, Allison N Rozanski, Peter Ebert, Glenn Hickey, Patrick Hasenfeld, Ashley D Sanders, Catherine Stober, Jan O Korbel, et al. Gaps and complex structurally variant loci in phased genome assemblies. Genome research, 33(4):496–510, 2023.

53. Peter Ebert, Peter A Audano, Qihui Zhu, Bernardo Rodriguez-Martin, David Porubsky, Marc Jan Bonder, Arvis Sulovari, Jana Ebler, Weichen Zhou, Rebecca Serra Mari, et al. Haplotype-resolved diverse human genomes and integrated analysis of structural variation. Science, 372(6537):eabf7117, 2021.

54. Peter Edge and Vikas Bansal. Longshot enables accurate variant calling in diploid genomes from single-molecule long read sequencing. Nat. Commun., 10(1):1–10, 2019.

55. Moritz Smolka, Luis F Paulin, Christopher M Grochowski, Dominic W Horner, Medhat Mahmoud, Sairam Behera, Ester Kalef-Ezra, Mira Gandhi, Karl Hong, Davut Pehlivan, et al. Detection of mosaic and population-level structural variants with sniffles2. Nature biotechnology, pages 1–10, 2024.

56. Marcel Martin, Murray Patterson, Shilpa Garg, Sarah O Fischer, Nadia Pisanti, Gunnar W Klau, Alexander Schöenhuth, and Tobias Marschall. Whatshap: fast and accurate read-based phasing. BioRxiv, page 085050, 2016.

57. Jyun-Hong Lin, Liang-Chi Chen, Shu-Chi Yu, and Yao-Ting Huang. Longphase: an ultra-fast chromosome-scale phasing algorithm for small and large variants. Bioinformatics, 38(7):1816–1822, 2022.

58. Wenlong Jia, Chang Xu, and Shuai Cheng Li. Resolving complex structures at oncovirus integration loci with conjugate graph. Briefings in Bioinformatics, 22(6):bbab359, 2021.

59. Chaohui Li, Lingxi Chen, Guangze Pan, and Wenqian Zhang. Deciphering complex breakage-fusion-bridge genome rearrangements with ambigram. Nature Communications, 14, 09 2023.

60. Justin M Zook, Jennifer McDaniel, Nathan D. Olson, Justin Wagner, Hemang Parikh, Haynes Heaton, Sean A. Irvine, and Len Trigg. An open resource for accurately benchmarking small variant and reference calls. Nature Biotechnology, 37:561–566, 2019.

61. Yukiteru Ono, Michiaki Hamada, and Kiyoshi Asai. Pbsim3: a simulator for all types of pacbio and ont long reads. NAR Genomics and Bioinformatics, 4(4):lqac092, 2022.

62. Andrea Gaedigk, Amy Turner, Robin E Everts, Stuart A Scott, Praful Aggarwal, Ulrich Broeckel, Gwendolyn A McMillin, Roberta Melis, Erin C Boone, Victoria M Pratt, et al. Characterization of reference materials for genetic testing of cyp2d6 alleles: a get-rm collaborative project. The Journal of Molecular Diagnostics, 21(6):1034–1052, 2019.

63. Esteban Arrieta-Bolaños, Diana Iraíz Hernández-Zaragoza, and Rodrigo Barquera. An hla map of the world: A comparison of hla frequencies in 200 worldwide populations reveals diverse patterns for class i and class ii. Frontiers in Genetics, 14:866407, 2023.

64. L DiAnne Bradford. Cyp2d6 allele frequency in european caucasians, asians, africans and their descendants. Pharmacogenomics, 3(2):229–243, 2002.

65. Glenys Thomson and Richard M Single. Conditional asymmetric linkage disequilibrium (ald): extending the biallelic r2 measure. Genetics, 198(1):321–331, 2014.

66. Kazutoyo Osoegawa, Steven J Mack, Matthew Prestegaard, and Marcelo A Fernández-Viña. Tools for building, analyzing and evaluating hla haplotypes from families. Human immunology, 80(9):633–643, 2019.

67. Nicholas R Pollock, Genelle F Harrison, and Paul J Norman. Immunogenomics of killer cell immunoglobulin-like receptor (kir) and hla class i: coevolution and consequences for human health. The Journal of Allergy and Clinical Immunology: In Practice, 10(7):1763–1775, 2022.

68. Nemr R. Finan R.R. Almawi, W.Y. et al. Hla-a, -b, -c, -drb1 and -dqb1 allele and haplotype frequencies in lebanese and their relatedness to neighboring and distant populations. BMC genomics, 23:456, 2022.

69. Leonardo M Amorim, Danillo G Augusto, Neda Nemat-Gorgani, Gonzalo Montero-Martin, Wesley M Marin, Hengameh Shams, Ravi Dandekar, Stacy Caillier, Peter Parham, Marcelo A Fernández-Viña, et al. High-resolution characterization of kir genes in a large north american cohort reveals novel details of structural and sequence diversity. Frontiers in Immunology, 12:674778, 2021.

70. Annika Niehrs and Marcus Altfeld. Regulation of nk-cell function by hla class ii. Frontiers in Cellular and Infection Microbiology, 10:55, 2020.

71. Gerd A Kullak-Ublick, Raul J Andrade, Michael Merz, Peter End, Andreas Benesic, Alexander L Gerbes, and Guruprasad P Aithal. Drug-induced liver injury: recent advances in diagnosis and risk assessment. Gut, 66(6):1154–1164, 2017.

72. Zeming Lin, Halil Akin, Roshan Rao, Brian Hie, Zhongkai Zhu, Wenting Lu, Nikita Smetanin, Robert Verkuil, Ori Kabeli, Yaniv Shmueli, Allan dos Santos Costa, Maryam Fazel-Zarandi, Tom Sercu, Salvatore Candido, and Alexander Rives. Evolutionary-scale prediction of atomic-level protein structure with a language model. Science, 379(6637):1123–1130, 2023.

73. Li C. Xue, João Pglm Rodrigues, Panagiotis L. Kastritis, Alexandre Mjj Bonvin, and Anna Vangone. Prodigy: a web server for predicting the binding affinity of protein–protein complexes. Bioinformatics, 32(23):3676–3678, 08 2016.

74. Caroline F. Thorn, Teri E. Klein, and Russ B. Altman. PharmGKB: The Pharmacogenomics Knowledge Base, pages 311–320. Humana Press, Totowa, NJ, 2013.

75. Michelle Whirl-Carrillo, Rachel Huddart, Li Gong, Katrin Sangkuhl, Caroline F. Thorn, Ryan Whaley, and Teri E. Klein. An evidence-based framework for evaluating pharmacogenomics knowledge for personalized medicine. Clinical Pharmacology & Therapeutics, 110(3):563–572, 2021.

76. Dominic J Barker, Giuseppe Maccari, Xenia Georgiou, Michael A Cooper, Paul Flicek, James Robinson, and Steven GE Marsh. The ipd-imgt/hla database. Nucleic acids research, 51(D1):D1053–D1060, 2023.

77. James Robinson, Jason A Halliwell, Hamish McWilliam, Rodrigo Lopez, and Steven GE Marsh. Ipd—the immuno polymorphism database. Nucleic acids research, 41(D1):D1234–D1240, 2012.

78. Véronique Giudicelli, Xavier Brochet, and Marie-Paule Lefranc. Imgt/v-quest: Imgt standardized analysis of the immunoglobulin (ig) and t cell receptor (tr) nucleotide sequences. Cold Spring Harbor Protocols, 2011(6):pdb–prot5633, 2011.

79. Andrea Gaedigk, Scott T Casey, Michelle Whirl-Carrillo, Neil A Miller, and Teri E Klein. Pharmvar: a global resource and repository for pharmacogene variation. Clinical pharmacology and therapeutics, 110(3):542, 2021.

80. Heng Li and Richard Durbin. Fast and accurate short read alignment with burrows–wheeler transform. bioinformatics, 25(14):1754–1760, 2009.

81. Heng Li. Minimap2: pairwise alignment for nucleotide sequences. Bioinformatics, 34(18):3094–3100, 2018.

82. Ryan Poplin, Pi-Chuan Chang, David Alexander, Scott Schwartz, Thomas Colthurst, Alexander Ku, Dan Newburger, Jojo Dijamco, Nam Nguyen, Pegah T Afshar, et al. A universal snp and small-indel variant caller using deep neural networks. Nature biotechnology, 36(10):983–987, 2018.

83. Stephen F Altschul, Thomas L Madden, Alejandro A Schäffer, Jinghui Zhang, Zheng Zhang, Webb Miller, and David J Lipman. Gapped blast and psi-blast: a new generation of protein database search programs. Nucleic Acids Res., 25(17):3389–3402, 1997.

84. Thomas Abeel, Thomas Van Parys, Yvan Saeys, James Galagan, and Yves Van de Peer. Genomeview: a next-generation genome browser. Nucleic Acids Research, 40(2):e12–e12, 2012.

85. Sam Kovaka, Aleksey V Zimin, Geo M Pertea, Roham Razaghi, Steven L Salzberg, and Mihaela Pertea. Transcriptome assembly from long-read rna-seq alignments with stringtie2. Genome biology, 20:1–13, 2019.

86. Fusheng Zhou, Hongzhi Cao, Xianbo Zuo, Tao Zhang, Xiaoguang Zhang, Xiaomin Liu, Ricong Xu, Gang Chen, Yuanwei Zhang, Xiaodong Zheng, et al. Deep sequencing of the mhc region in the chinese population contributes to studies of complex disease. Nature genetics, 48(7):740–746, 2016.

87. Donald F Conrad, Jonathan E M Keebler, Mark A DePristo, Sarah J Lindsay, Yujun Zhang, Ferran Casals, Youssef Idaghdour, Chris L Hartl, Carlos Torroja, Kiran V Garimella, Martine Zilversmit, Reed Cartwright, Guy A Rouleau, Mark Daly, Eric A Stone, Matthew E Hurles, Philip Awadalla, and the 1000 Genomes Project. Variation in genome-wide mutation rates within and between human families. Nature Genetics, 43:712–714, 2011.

88. Jared C. Roach, Gustavo Glusman, Arian F. A. Smit, Chad D. Huff, Robert Hubley, Paul T. Shannon, Lee Rowen, Krishna P. Pant, Nathan Goodman, Michael Bamshad, Jay Shendure, Radoje Drmanac, Lynn B. Jorde, Leroy Hood, and David J. Galas. Analysis of genetic inheritance in a family quartet by whole-genome sequencing. Science, 328(5978):636–639, 2010.

89. Michael Lynch. Rate, molecular spectrum, and consequences of human mutation. Proceedings of the National Academy of Sciences, 107(3):961–968, 2010.

90. John N Feder. The hereditary hemochromatosis gene (hfe). Immunologic Research, 20, 1999.

91. Ping Song, Qiang Zhao, and Ming-Hui Zou. Targeting senescent cells to attenuate cardiovascular disease progression. Ageing Research Reviews, 60:101072, 2020.

92. Woong-Kyung Suh, Myrna F. Cohen-Doyle, Klaus Fruh, Kena Wang, Per A. Peterson, and David B. Williams. Interaction of mhc class i molecules with the transporter associated with antigen processing. Science, 264(5163):1322–1326, 1994.

