## Supplementary Material for "SpecImmune accurately genotypes diverse immune-related gene families using long-read data"

### Supplemental Information

#### Supplemental Note

##### Note S1 Computational resource evaluation of SpecImmune

SpecImmune represents a swift method for immune-related gene typing. In assessing the computational resource utilization of SpecImmune, we undertook HLA, KIR, IG+TCR, and CYP typing individually utilizing the HPRC HiFi dataset, which encompassed 50 PacBio HiFi WGS samples. Reads from each gene family were isolated, and SpecImmune was independently executed for each gene family. Subsequently, we documented the wall-clock time, CPU time, and peak memory usage for each execution. Initially, we juxtaposed the computational resource utilization of SpecImmune against SpecHLA and HLA*LA for HLA typing. We performed the three methods with default parameters using 20 threads. Given the variance in the number of supported gene loci among these three HLA typing methods, we normalized the wall-clock time and CPU time concerning the number of supported gene loci. Notably, the normalized wall-clock time was found to be the shortest for SpecImmune (Figure S3). The normalized CPU time and peak memory usage exhibited similarities between SpecImmune and SpecHLA. In contrast, HLA*LA exhibited significantly higher peak memory consumption and normalized CPU time than the other two methods. On average, SpecImmune required around 10.85G of Peak RAM, 2.22 hours of CPU time, and 0.45 hours of wall-clock time per sample.

In KIR typing, the average CPU time and peak memory usage are 8.78 hours and 0.94 GB, respectively (Figure S4). SpecImmune facilitates simultaneous IG and TCR typing. For IG+TCR typing, SpecImmune exhibits an average CPU time of 8.72 hours and a peak memory usage of 7.33 GB (Figure S4). In CYP2D6 typing, the average CPU time and peak memory usage are 0.49 hours and 11.85 GB, respectively (Figure S4). SpecImmune demonstrates efficient memory and time management in typing immune-related genes, making it suitable for execution on standard personal computers. This capability enables SpecImmune to provide real-time gene typing results in tandem with contemporary sequencing technologies, enhancing its utility for clinical applications.

##### Note S2 HLA and KIR allele frequency exhibit variation across loci

Common, low-frequency, and rare HLA allele frequencies exhibit variation across gene loci. We categorized the alleles based on their frequencies within the population (Methods); 9.4% (187), 23.1% (459), and 67.6% (1345) of these identified alleles were classified as common, low-frequency, and rare, respectively (Figure S10). We then analyzed the frequency distribution of these three groups within each gene locus. Notably, none of the *HLA-B* and *HLA-DRB1* alleles were classified as common, while all *HLA-DPB2*, *HLA-V*, *HLA-S*, and *HLA-Y* alleles fell into this category. Among the genes, *MICB* exhibited the highest rare allele frequency at 87.2%, followed by *HLA-DPB1* at 84.8% and *HLA-A* at 83.6%.

Furthermore, the occurrence of rare alleles varies significantly among the KIR loci. We examined the frequency distribution of common, low-frequency, and rare alleles for each locus (Figure S10). *KIR2DL2* and *KIR2DL5B* exhibit no rare alleles, with relatively low frequencies of rare alleles observed in *KIR2DL3*, *KIR2DS1*, and *KIR2DS4*. In contrast, the majority of alleles in *KIR2DL1* and *KIR3DL1* are classified as rare.

##### Note S3 HLA and KIR allele diversity exhibit variation across loci

HLA allele diversity exhibits variability across gene loci. The gene diversity within each of the 26 populations was quantified by the Shannon Diversity Index (Methods). Notably, there was significant variation in diversity across the HLA loci (Figure S11). *HLA-DRB1* displayed the highest mean diversity among the populations, followed by *HLA-DPB1*, *HLA-B*, *HLA-A*, *HLA-C*, and *HLA-DQB1*. The notable diversity of these genes is consistent with previous findings in the Han Chinese population [1]. The hierarchy of diversity among *HLA-B*, *HLA-A*, and *HLA-C* conforms to previous research, suggesting that *HLA-B* is the oldest and most diverse locus, while *HLA-C* has evolved more recently [2]. Conversely, HLA-Y had the lowest mean diversity, with *HLA-U*, *HFE*, and *HLA-L* following suit. Furthermore, there were no significant differences observed in gene diversity between populations.

The Shannon diversity exhibits significant variation across the KIR loci. We computed the frequency of each allele and determined the Shannon diversity at each KIR locus (Methods). *KIR2DL1* demonstrated the highest Shannon diversity, followed by *KIR3DP1* (Figure S12). In contrast, *KIR2DL2* displayed the lowest Shannon diversity, with *KIR2DS4* following suit. In certain loci, a small number of alleles exhibited notably high frequencies. For instance, the allele KIR3DS1*0130101 (23.2%) emerged as the most frequent among all KIR alleles, closely trailed by KIR2DS1*0020103 (22.6%).

##### Note S4 Heterozygosity levels vary significantly across IG/TCR gene loci

The heterozygosity levels vary significantly across IG/TCR gene loci. A locus was considered heterozygous if two different alleles were present and homologous otherwise within a sample. We calculated the heterozygous frequencies for each locus across all samples (Figure S13a-b). Interestingly, 57.4% (220 out of 402) of the loci exhibit a 0% heterozygous frequency. On the other hand, *IGHV1-69* exhibited the highest heterozygous frequency at 62.4%, followed by *IGHV3-48* at 58.4%, and *TRAV12-2* at 56.8%.

##### Note S5 The discovery of DNMs

In the six parent-offspring trios we studied, the DNMs rate in the MHC region averaged $1.42 \times 10^{-4}$, which is significantly higher than the genome-wide DNM rate of $1.1 \times 10^{-8}$ to $3 \times 10^{-8}$ per generation, as reported in previous studies [3, 4, 5, 6]. This substantial increase suggests that an elevated mutation rate partly drives the high genetic diversity in the MHC region. The MHC, known for its polymorphism, is under strong evolutionary pressures, such as balancing selection and pathogen-driven selection, which likely promote both the retention of diverse alleles and higher mutational input [7, 8]. Additionally, unique genomic features of the MHC, such as increased recombination rates or distinct DNA repair mechanisms, may contribute to this elevated rate, further explaining the pronounced divergence observed in this region [9].

### References


[1] Fusheng Zhou, Hongzhi Cao, Xianbo Zuo, Tao Zhang, Xiaoguang Zhang, Xiaomin Liu, Ricong Xu, Gang Chen, Yuanwei Zhang, Xiaodong Zheng, et al. Deep sequencing of the mhc region in the chinese population contributes to studies of complex disease. *Nature genetics*, 48(7):740–746, 2016.

[2] Diego Chowell, Chirag Krishna, Federica Pierini, Vladimir Makarov, Naiyer A Rizvi, Fengshen Kuo, Luc GT Morris, Nadeem Riaz, Tobias L Lenz, and Timothy A Chan. Evolutionary divergence of hla class i genotype impacts efficacy of cancer immunotherapy. *Nature medicine*, 25(11):1715–1720, 2019.

[3] Jared C. Roach, Gustavo Glusman, Arian F. A. Smit, Chad D. Huff, Robert Hubley, Paul T. Shannon, Lee Rowen, Krishna P. Pant, Nathan Goodman, Michael Bamshad, Jay Shendure, Radoje Drmanac, Lynn B. Jorde, Leroy Hood, and David J. Galas. Analysis of genetic inheritance in a family quartet by whole-genome sequencing. *Science*, 328(5978):636–639, 2010.

[4] Donald F Conrad, Jonathan E M Keebler, Mark A DePristo, Sarah J Lindsay, Yujun Zhang, Ferran Casals, Youssef Idaghdour, Chris L Hartl, Carlos Torroja, Kiran V Garimella, Martine Zilversmit, Reed Cartwright, Guy A Rouleau, Mark Daly, Eric A Stone, Matthew E Hurles, Philip Awadalla, and the 1000 Genomes Project. Variation in genome-wide mutation rates within and between human families. *Nature Genetics*, 43:712–714, 2011.

[5] Michael W Nachman and Susan L Crowell. Estimate of the mutation rate per nucleotide in humans. *Genetics*, 156(1):297–304, 09 2000.

[6] Michael Lynch. Rate, molecular spectrum, and consequences of human mutation. *Proceedings of the National Academy of Sciences*, 107(3):961–968, 2010.

[7] Pierre-Antoine Gourraud, Pouya Khankhanian, Nezih Cereb, Soo Young Yang, Michael Feolo, Martin Maiers, John D. Rioux, Stephen Hauser, and Jorge Oksenberg. Hla diversity in the 1000 genomes dataset. *PLOS ONE*, 9(7):1–8, 07 2014.

[8] Wayne K. Potts and Edward K. Wakeland. Evolution of diversity at the major histocompatibility complex. *Trends in Ecology Evolution*, 5:0169–5347, 06 1990.

[9] Robert E. Hickson and Rebecca L. Cann. Mhc allelic diversity and modern human origins. *Journal of Molecular Evolution*, 45:589–598, 12 1997.

**Algorithm 1** Iterative Haplotype Reconstruction

**Input:**
Aligned reads $R$, personalized reference alleles $P$, window size for masking low-depth regions $w$ (default $= 20$ bp), depth threshold $\tau$ (default $= 5$).

**Output:**
Reconstructed personalized diploid haplotype sequences $H = \{H_1, H_2\}$.

```
Initialize haplotypes: H ← ∅
Initialize variant identification flag: variants_identified ← True
while variants_identified = True do
    (a) Realign reads:
    R′ ← Realign(R, H or P)
    (b) Variant calling:
    SNVs ← CallSNVs(R′)
    SV_breakpoints ← CallSVBreakpoints(R′)
    (c) Variant phasing:
    Phased_SNVs ← PhaseSNVs(SNVs, R′)
    Phased_SNVs_and_SVs ← JointPhase(Phased_SNVs, SV_breakpoints)
    (d) Consensus sequence reconstruction:
    Consensus ← GenerateConsensus(Phased_SNVs_and_SVs, P)
    (e) Segmentation:
    S = {s_1, s_2, ..., s_n} ← SegmentConsensus(Consensus, SV_breakpoints)
    (f) Copy number estimation:
    for s_i ∈ S do
        c_i ← ⌊ d_i / ((1/n) Σ_{j=1}^{n} d_j) ⌋
    end for
    (g) Graph construction and matching:
    Construct bipartite graph G(S, E) with edge weights:

        w(s_j, s_i) = Θ(s_j, s_i)   (spanning read count between s_j and s_i)

    H ← PerformConjugateMatching(G)
    (h) Mask low-depth regions:
    H ← MaskLowDepthRegions(H, R′, w, τ)
    (i) Variant identification:
    New_Variants ← IdentifyNewVariants(H, R′)
    variants_identified ← (New_Variants ≠ ∅)
end while
Return: H = {H_1, H_2}
```

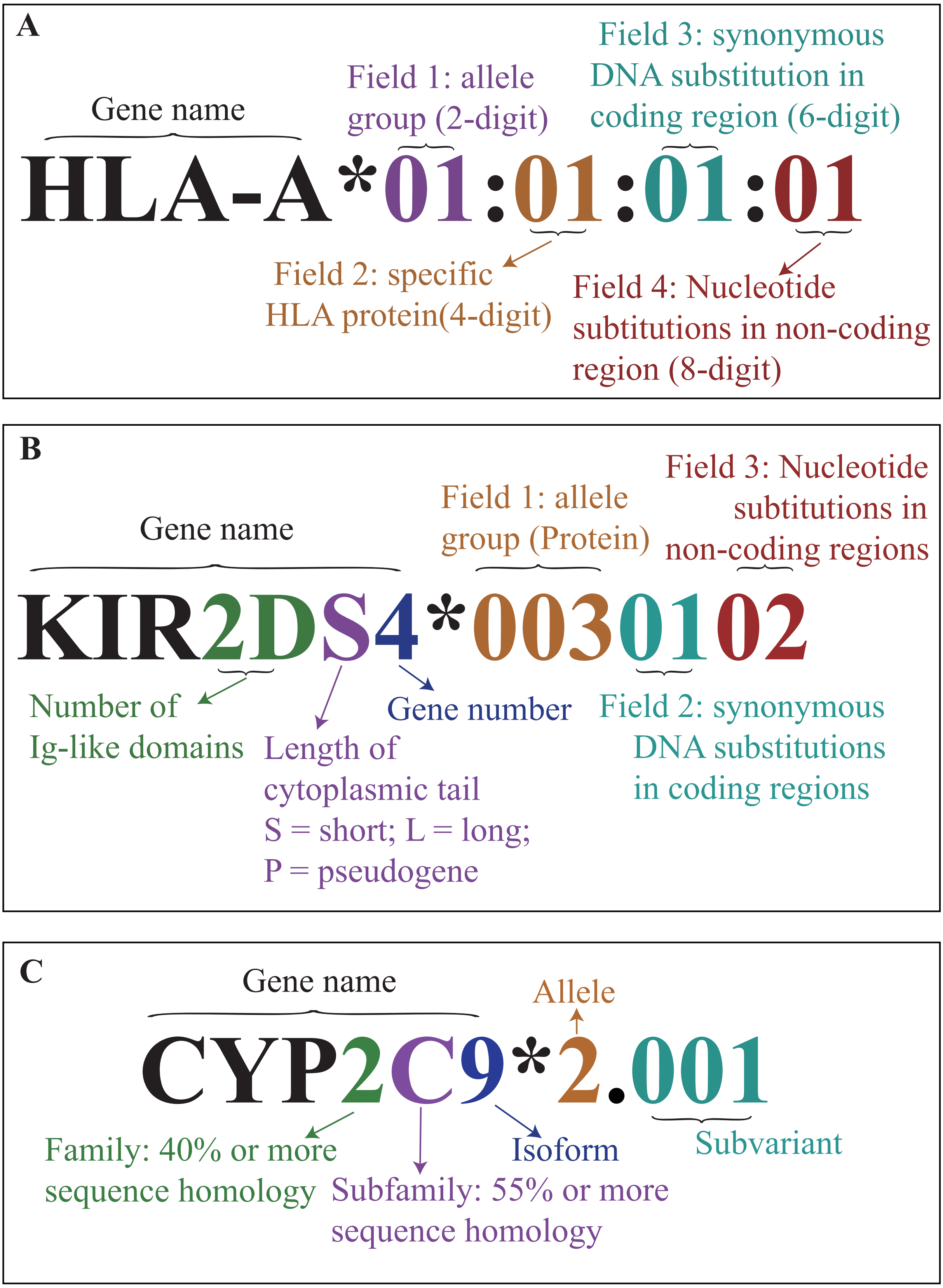


**Figure S1: Nomenclature description of HLA/KIR/CYP alleles.**
(A) HLA, (B) KIR, and (C) CYP nomenclature with different allele resolutions.

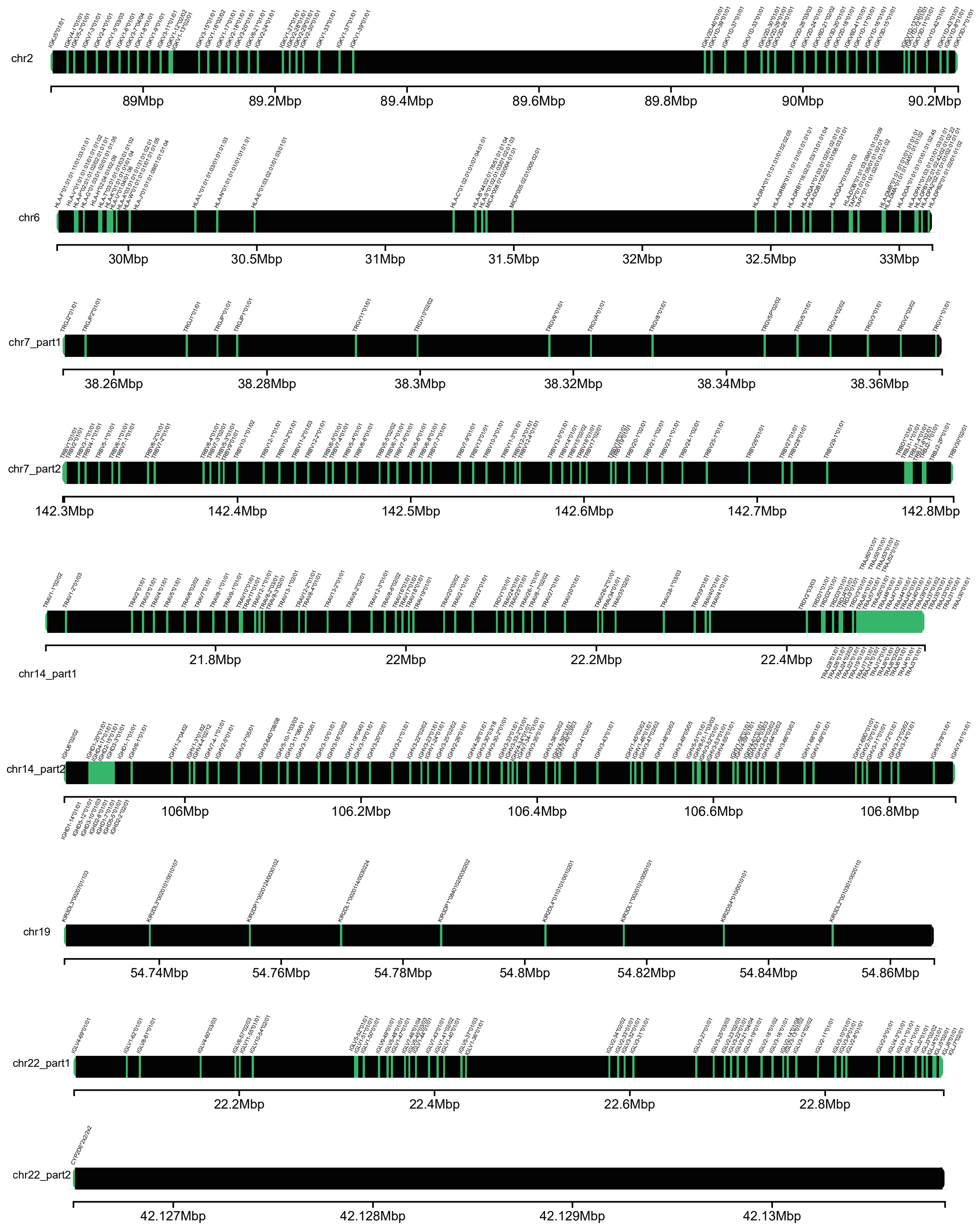


**Figure S2: Summary of SpecImmune typing results for HG00377 in 1kGP.** Visualization includes only loci on the main chromosome of hg38.

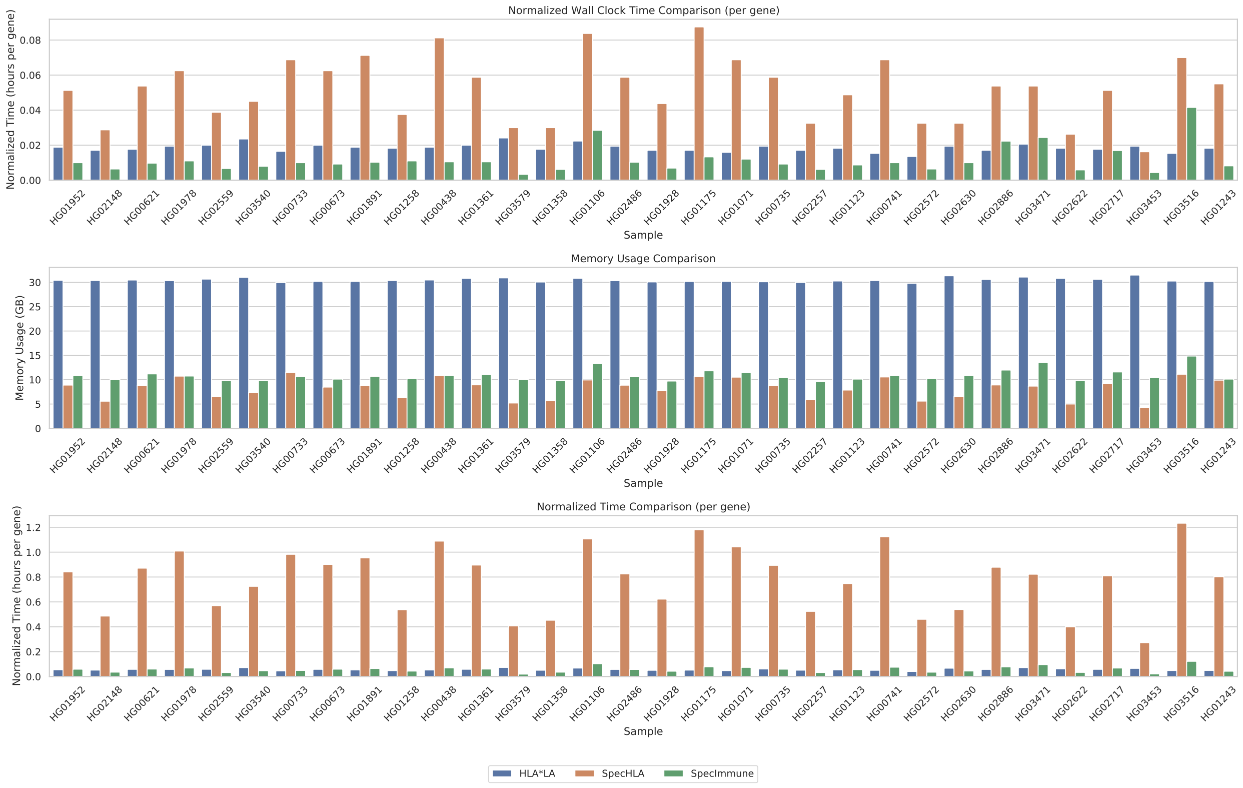


**Figure S3: Computational resource evaluation of HLA*LA, SpecHLA and SpecImmune for HLA typing in HPRC HiFi samples.**

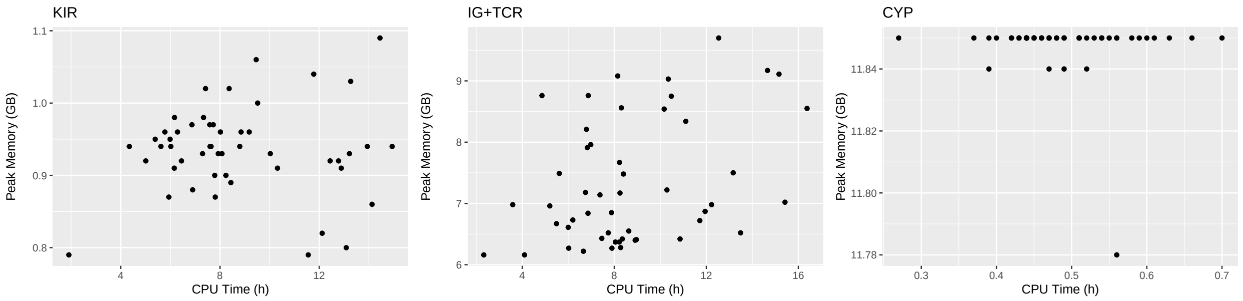


**Figure S4: Computational resource evaluation of SpecImmune for KIR, IG/TCR, and CYP typing on HPRC HiFi samples.**

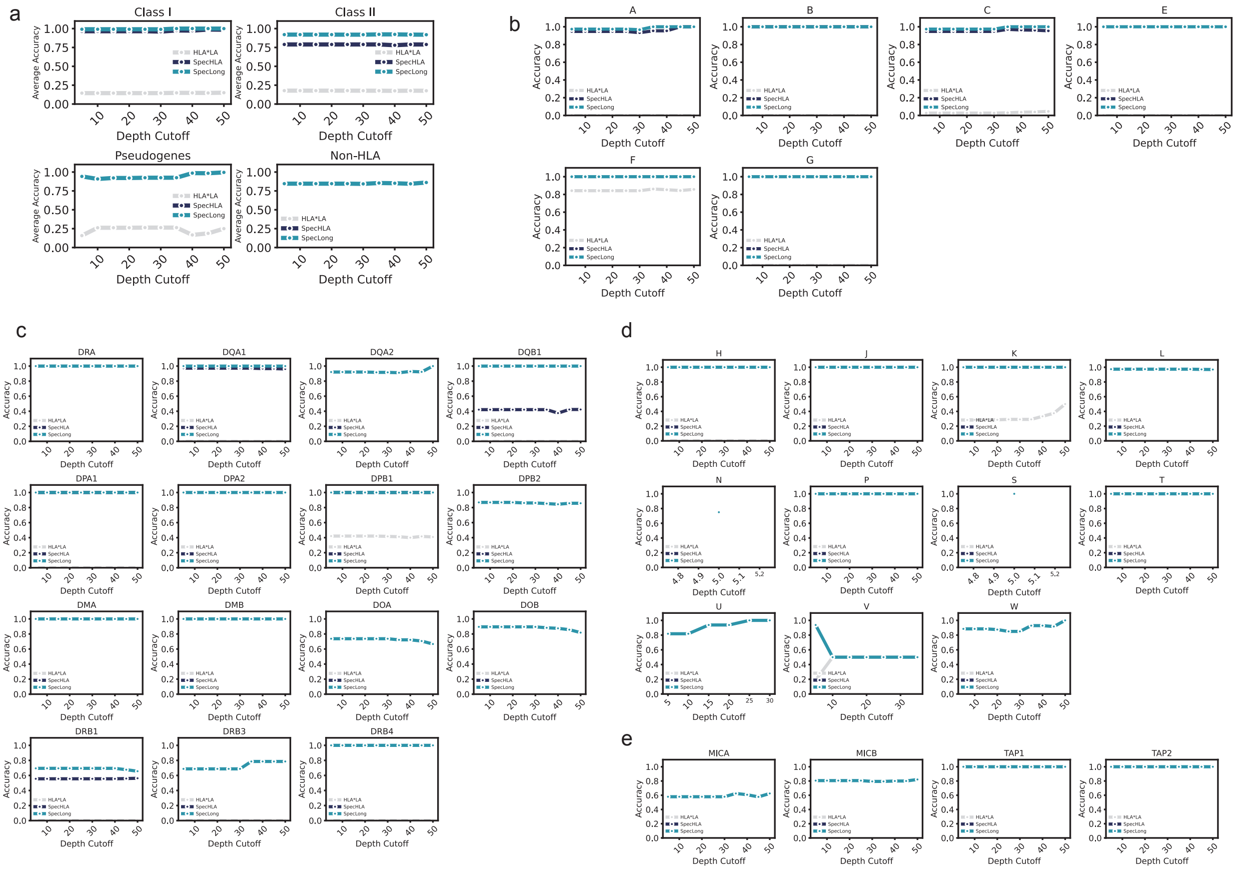


**Figure S5: Performance of HLA*LA, SpecHLA, and SpecImmune on HGSVC CLR dataset.** (a) Accuracy of HLA*LA, SpecHLA, and SpecImmune of 4 HLA gene classes. (b) Accuracy of HLA*LA, SpecHLA, and SpecImmune of HLA class I genes. (c) Accuracy of HLA*LA, SpecHLA, and SpecImmune of HLA class II genes. (d) Accuracy of HLA*LA, SpecHLA, and SpecImmune of HLA Pseudogenes genes. (e) Accuracy of HLA*LA, SpecHLA, and SpecImmune of Non-HLA genes.

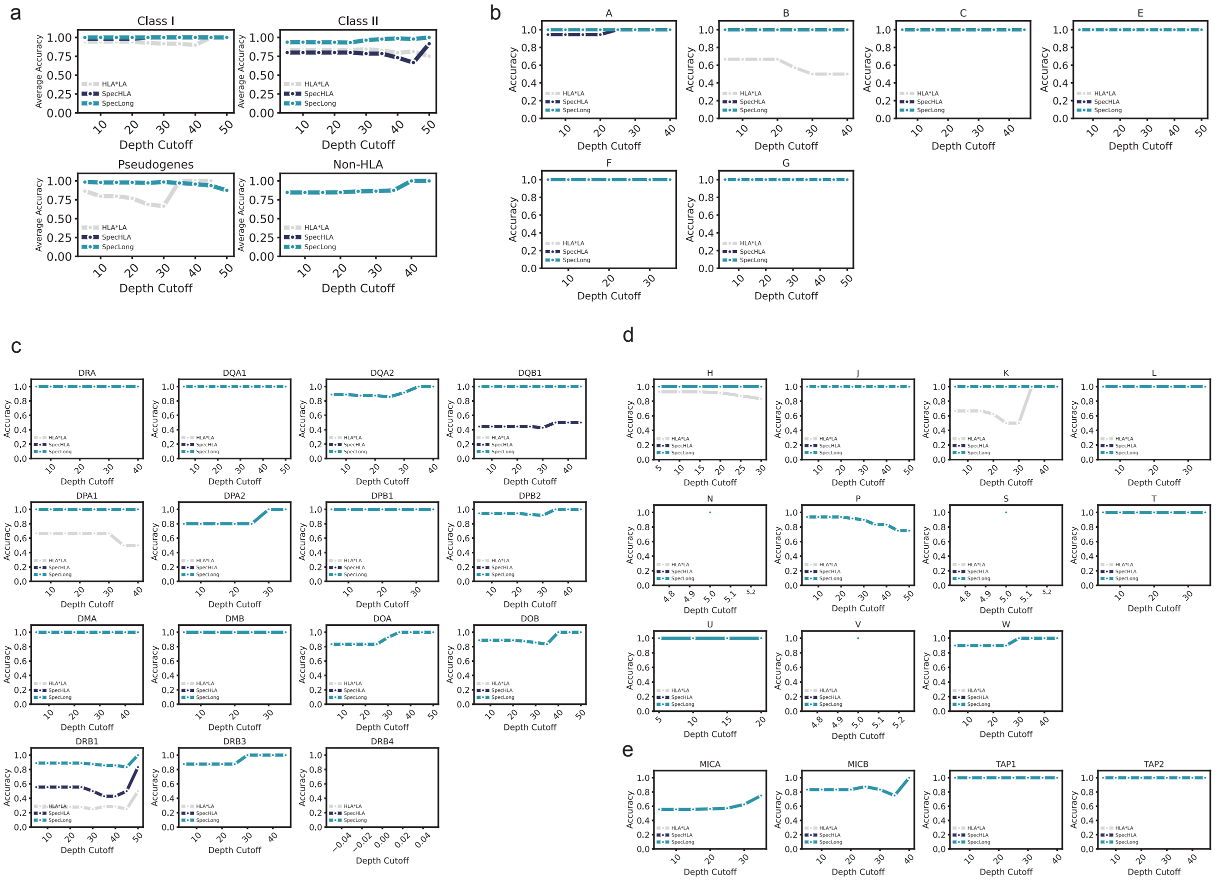


**Figure S6: Performance of HLA*LA, SpecHLA, and SpecImmune on HGSVC HiFi dataset.**
(a) Accuracy of HLA*LA, SpecHLA, and SpecImmune of 4 HLA gene classes. (b) Accuracy of HLA*LA, SpecHLA, and SpecImmune of HLA class I genes. (c) Accuracy of HLA*LA, SpecHLA, and SpecImmune of HLA class II genes. (d) Accuracy of HLA*LA, SpecHLA, and SpecImmune of HLA Pseudogenes genes. (e) Accuracy of HLA*LA, SpecHLA, and SpecImmune of Non-HLA genes.

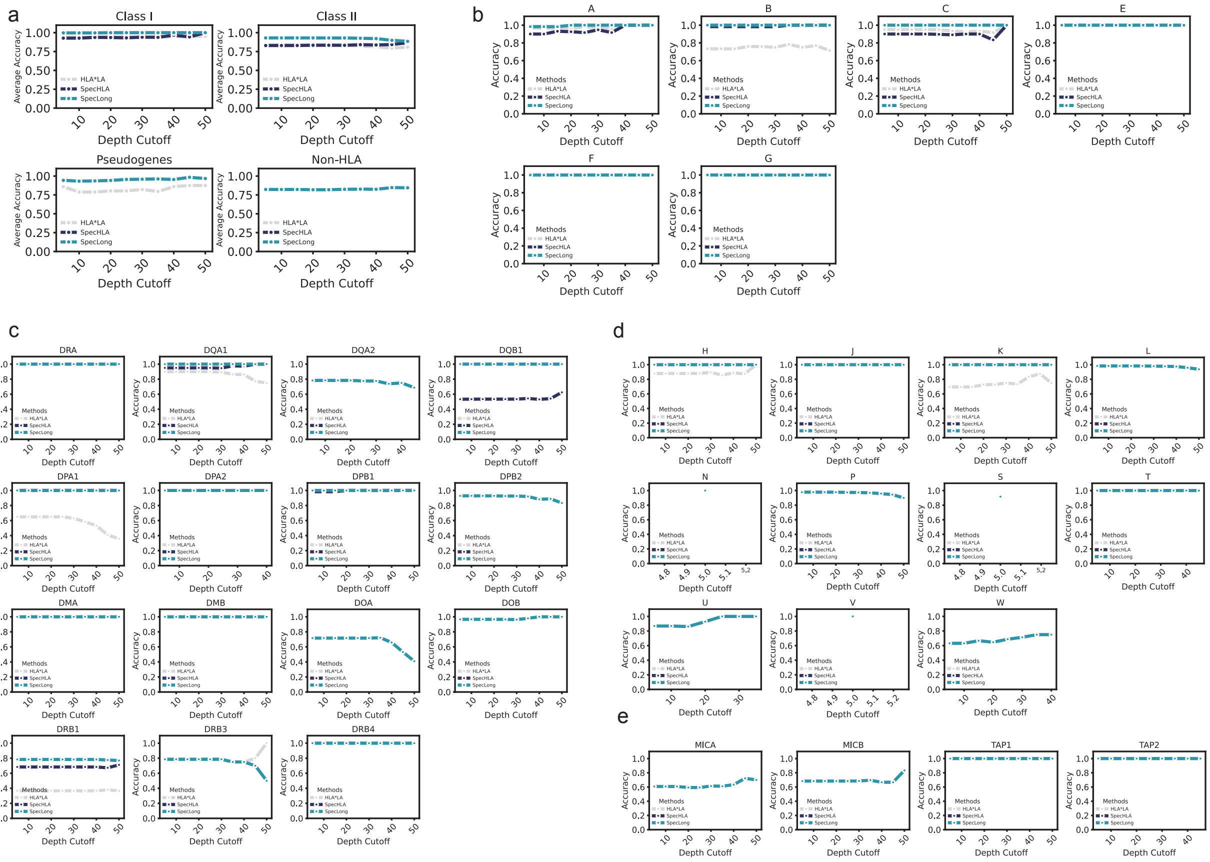


**Figure S7: Performance of HLA*LA, SpecHLA, and SpecImmune on HPRC HiFi dataset.** (a) Accuracy of HLA*LA, SpecHLA, and SpecImmune of 4 HLA gene classes. (b) Accuracy of HLA*LA, SpecHLA, and SpecImmune of HLA class I genes. (c) Accuracy of HLA*LA, SpecHLA, and SpecImmune of HLA class II genes. (d) Accuracy of HLA*LA, SpecHLA, and SpecImmune of HLA Pseudogenes genes. (e) Accuracy of HLA*LA, SpecHLA, and SpecImmune of Non-HLA genes.

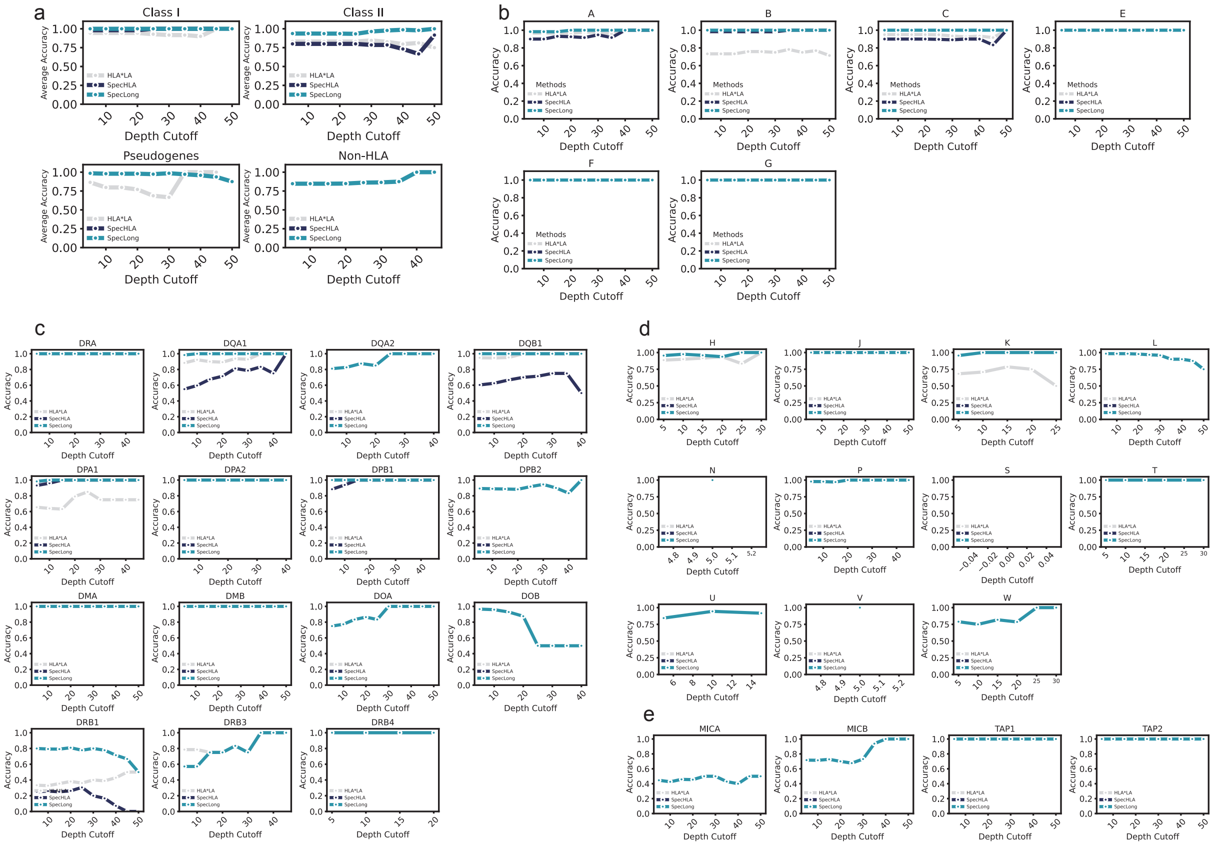


**Figure S8: Performance of HLA*LA, SpecHLA, and SpecImmune on HPRC ONT dataset.**
(a) Accuracy of HLA*LA, SpecHLA, and SpecImmune of 4 HLA gene classes. (b) Accuracy of HLA*LA, SpecHLA, and SpecImmune of HLA class I genes. (c) Accuracy of HLA*LA, SpecHLA, and SpecImmune of HLA class II genes. (d) Accuracy of HLA*LA, SpecHLA, and SpecImmune of HLA Pseudogenes genes. (e) Accuracy of HLA*LA, SpecHLA, and SpecImmune of Non-HLA genes.

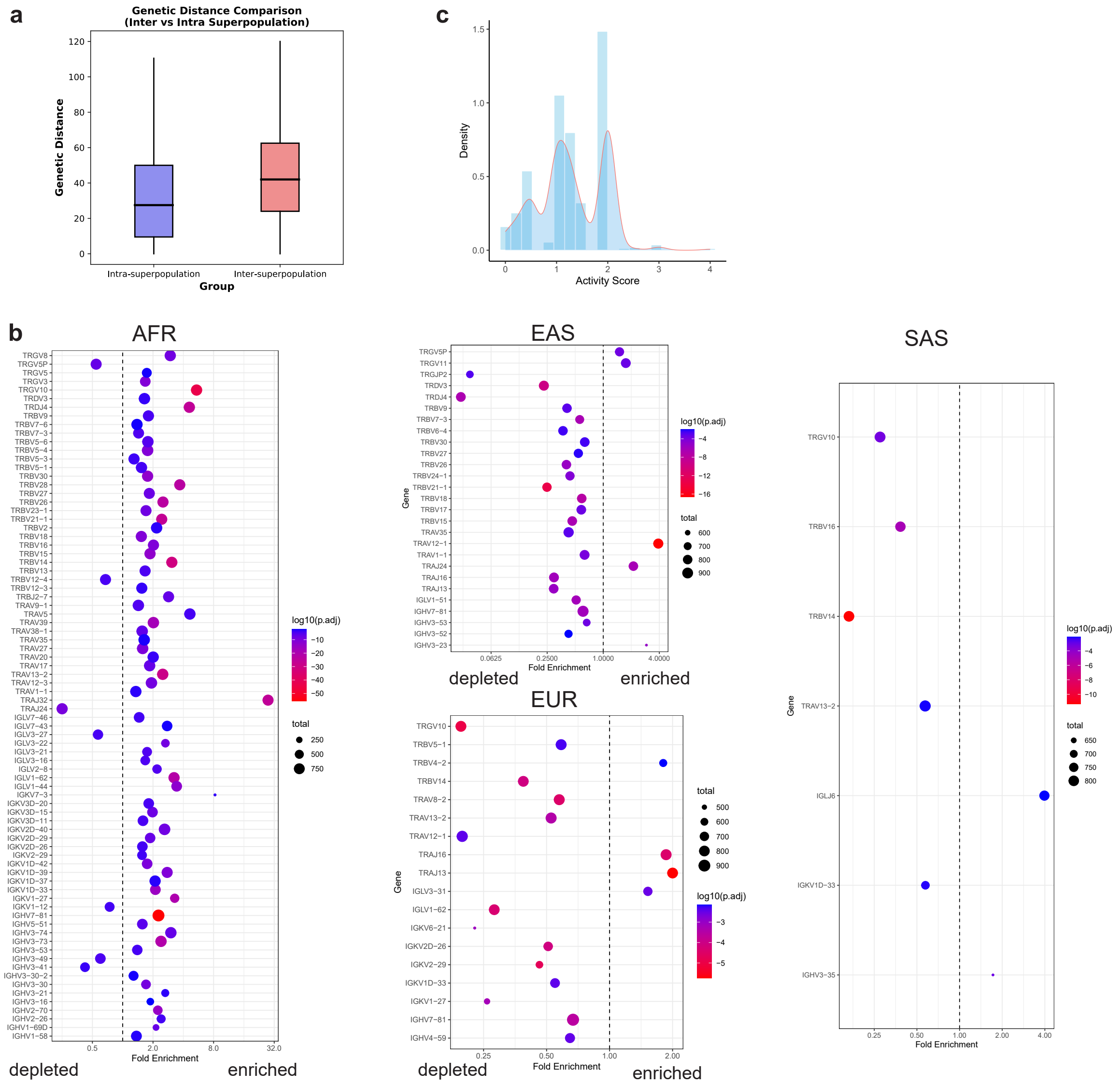


**Figure S9: Landscape of immune-related gene alleles in 1kGP population.**
(a) Comparison of HLA genetic distance within the same super populations and between different super populations. (b) Genes with enriched and depleted heterozygous variants in the populations. (c) Distribution of *CYP2D6* activity scores across all samples.

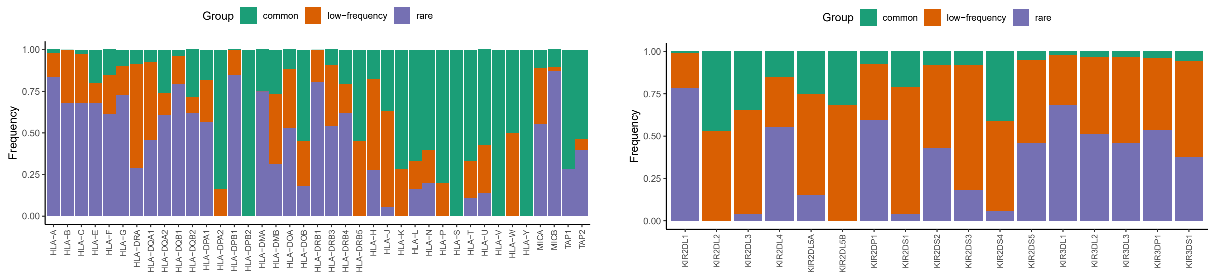


**Figure S10: Frequencies of common, low-frequency, and rare alleles at each HLA and KIR locus**

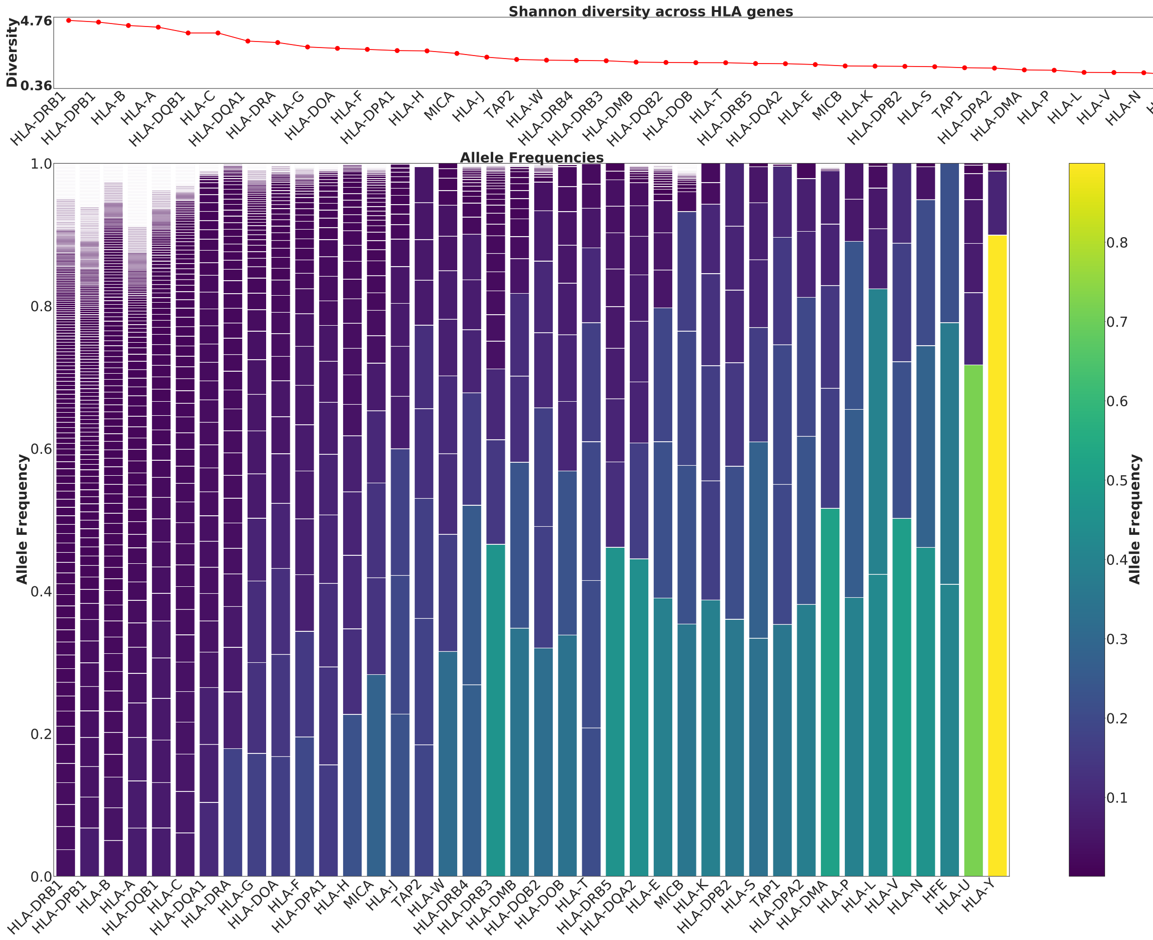


**Figure S11: Allelic diversity at each HLA locus.**

Bottom barplot indicates the frequency of each allele at each locus.

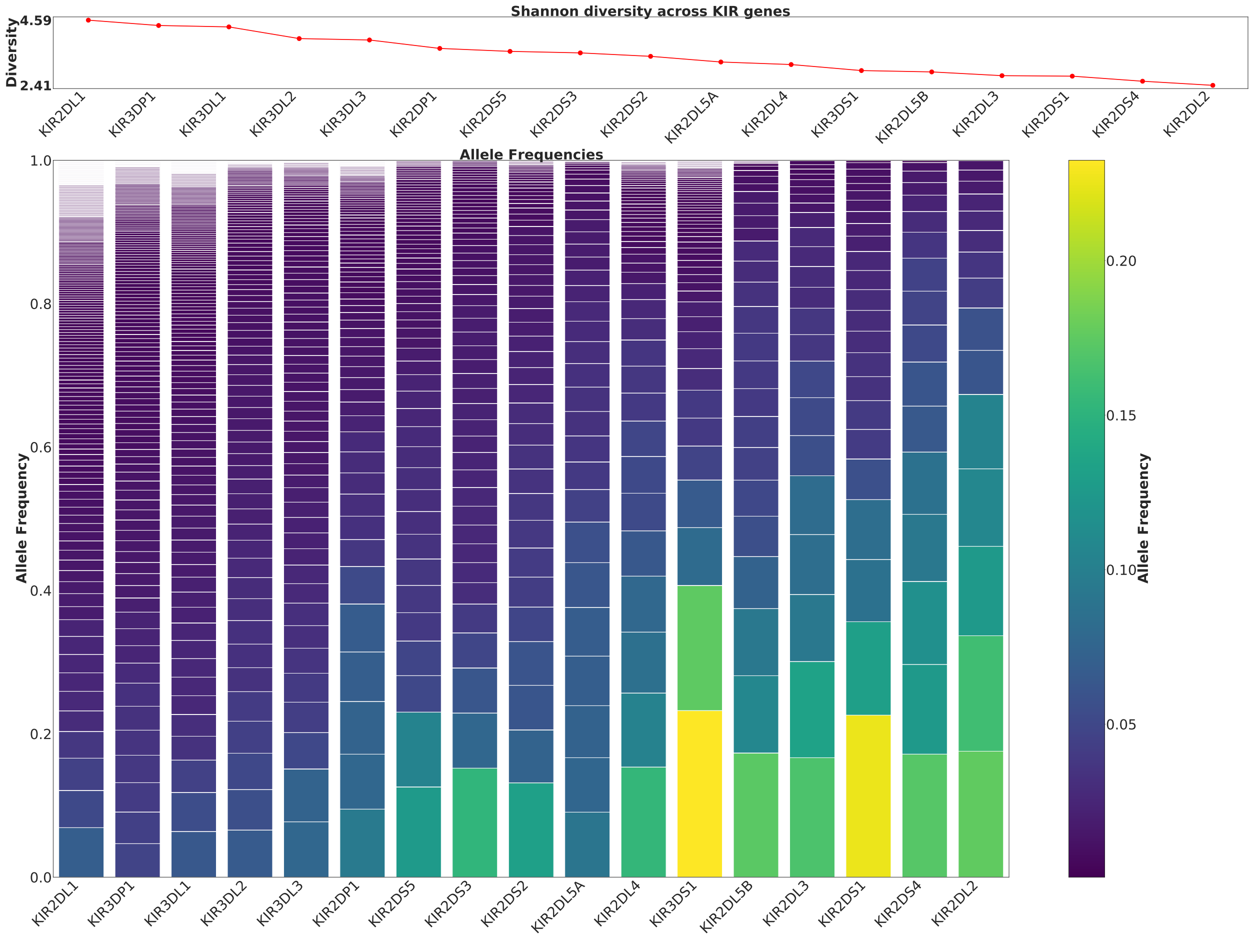


**Figure S12: Allelic diversity at each KIR locus.**

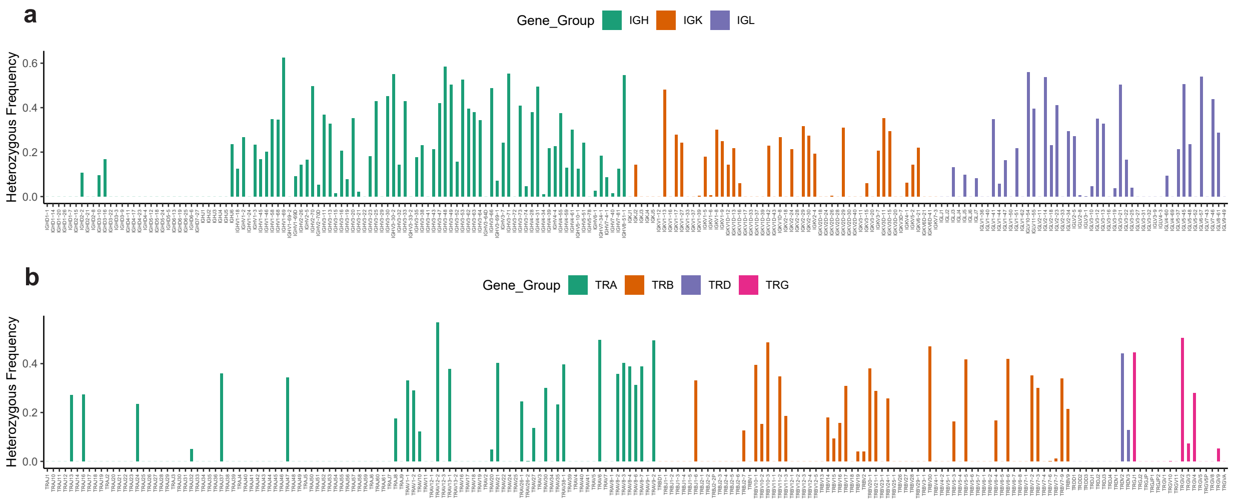


**Figure S13: (a-b) Heterozygous frequencies at each IG (a) and TCR (b) locus.**

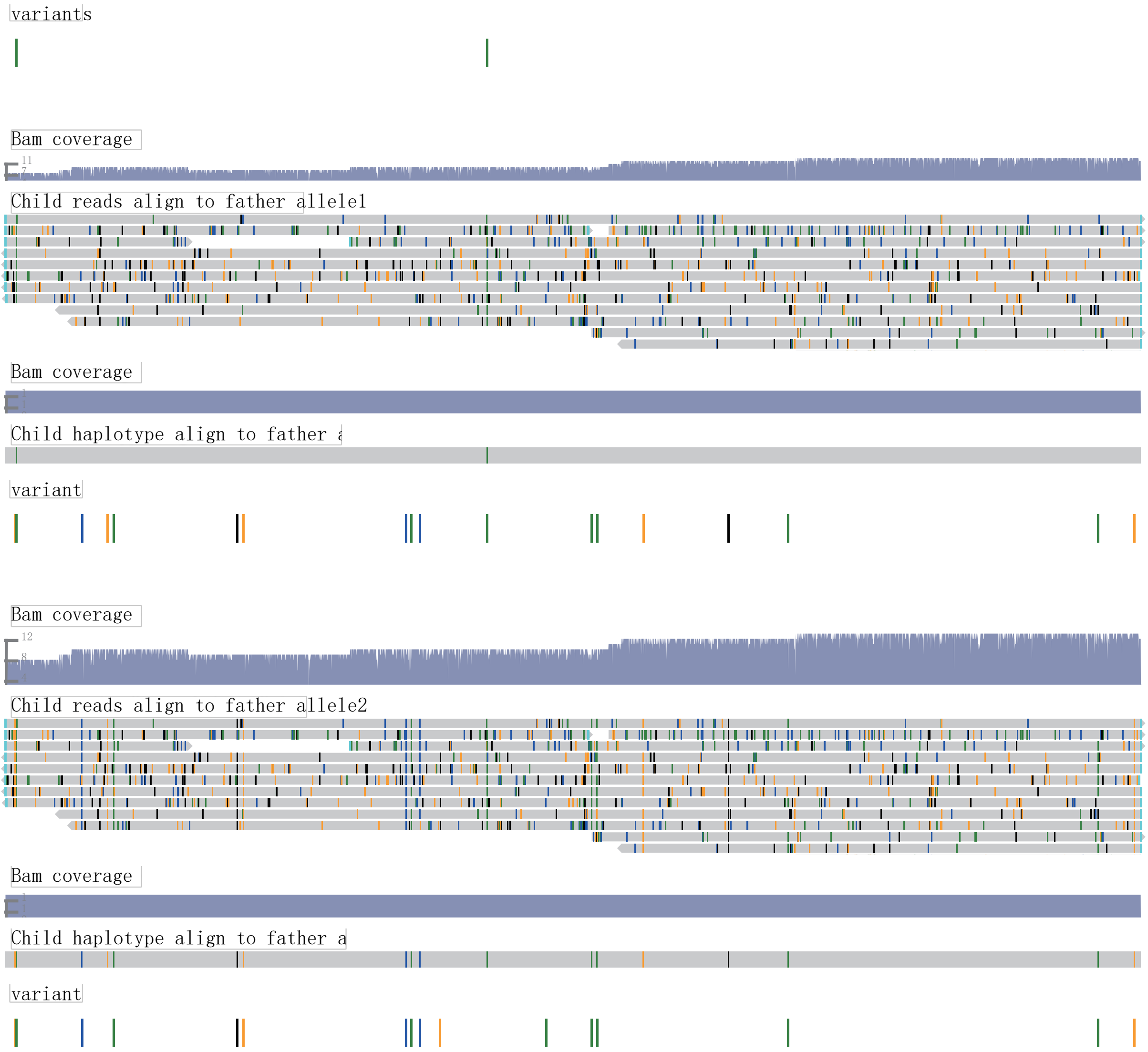


**Figure S14: Illustration of novel variants detected by SpecImmune.** Novel variants have been detected at the *HLA-DOB* locus in the family trio SH006, where the child exhibits homozygosity. The figure illustrates the child's haplotype aligned with those of the father. Variants are color-coded within the alignment for clarity.

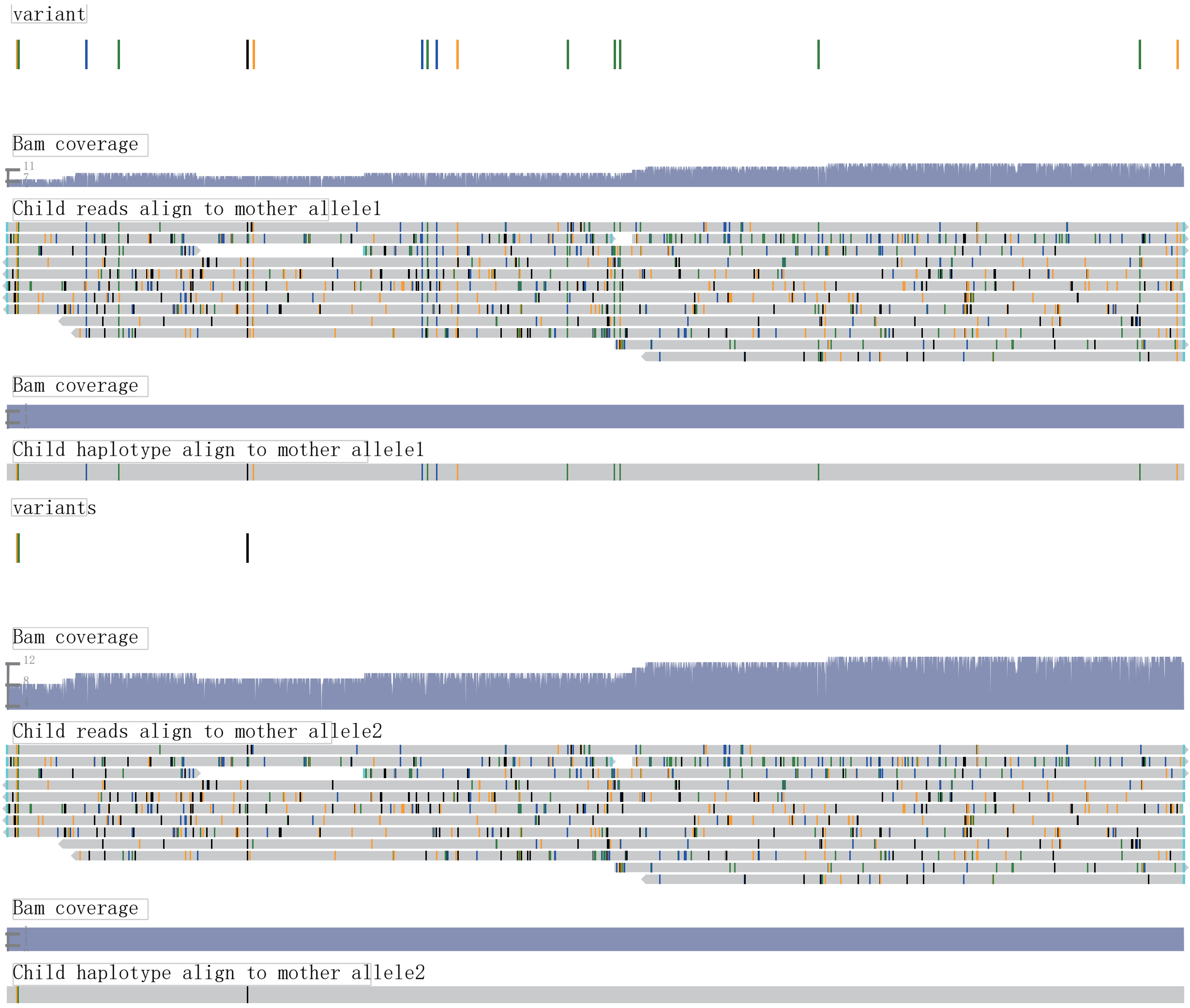


**Figure S15: Illustration of novel variants detected by SpecImmune.** Novel variants have been detected at the *HLA-DOB* locus in the family trio SH006, where the child exhibits homozygosity. The figure illustrates the child's haplotype aligned with those of the mother. Variants are color-coded within the alignment for clarity.

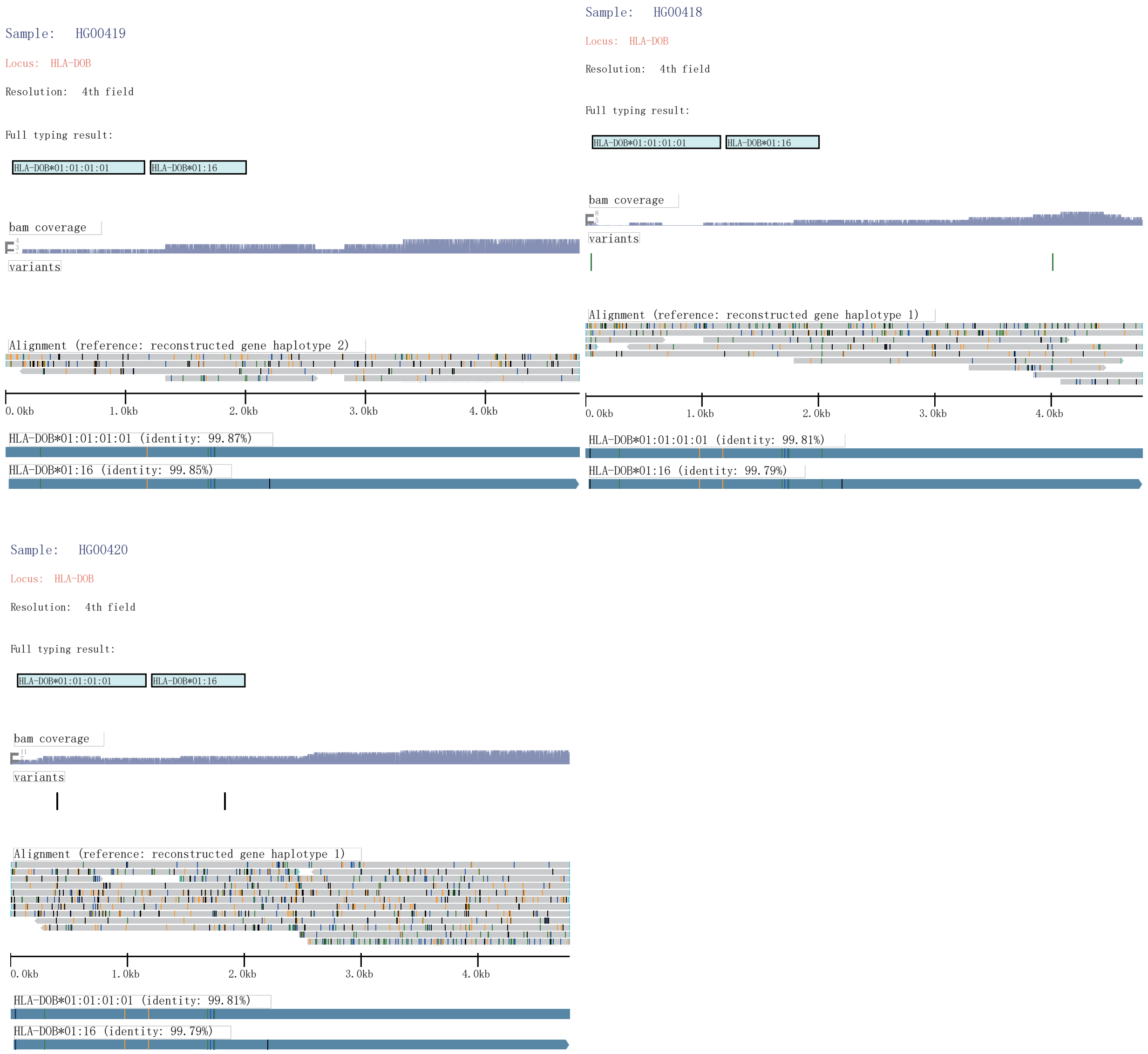


**Figure S16: Illustration of inherited alleles detected by SpecImmune.** Allele sequences and supporting reads detected at the *HLA-DOB* locus in the family trio SH006, where the child exhibits homozygosity. Variants are color-coded within the alignment for clarity. HG00420 is child, HG00418 is father, and HG00419 is mother.

**Table S1** Commands used for SpecHLA, HLA*LA, and SpecImmune in evaluation

| Software | Command | Gene Family |
|---|---|---|
| **SpecHLA** | `python3 SpecHLA/script/long_read_typing.py -r $fastq -j 20 -n $sample_id -o $outdir -y $datatype` | HLA |
| **HLA*LA** | `HLA-LA/src/HLA-LA.pl -BAM $sample_bam -graph PRG_MHC_GRCh38_withIMGT -sampleID $sample_id -maxThreads 20 -longReads $datatype -samtools_T $REFERENCE_GENOME -workingDir $outdir` | HLA |
| **SpecImmune** | `python3 SpecImmune/main.py -r $fastq -j 20 -i HLA -n $sample_id -o $outdir -db SpecImmune/db -y $datatype -align_method_1 minimap2` | HLA |
| **SpecImmune** | `python3 SpecImmune/main.py -r $fastq -j 20 -i KIR -n $sample_id -o $outdir -db SpecImmune/db -y $datatype -align_method_1 minimap2 -hete_p 0.2` | KIR |
| **SpecImmune** | `python3 SpecImmune/main.py -r $fastq -j 20 -i CYP -n $sample_id -o $outdir -db SpecImmune/db -y $datatype` | CYP |
| **SpecImmune** | `python3 SpecImmune/main.py -r $fastq -j 20 -i IG_TR -n $sample_id -o $outdir -db SpecImmune/db -y $datatype -hg38 $no_alt_ref` | IG and TCR |

**Table S2** Trio information in 1kGP

| Trio ID | Child | Father | Mother |
|---|---|---|---|
| 2418 | NA19828 | NA19818 | NA19819 |
| CLM16 | HG01258 | HG01256 | HG01257 |
| 1463-Paternal | NA12877 | NA12889 | NA12890 |
| 1463-Maternal | NA12878 | NA12891 | NA12892 |
| SH006 | HG00420 | HG00418 | HG00419 |
| Y077 | NA19129 | NA19128 | NA19127 |
